## Supplemental Material for "Diclofenac sensitizes multi-drug resistant *Acinetobacter baumannii* to colistin"

**Table S1: Strains used in this study.**

| | | |
|---|---|---|
| 17978 | *A. baumannii* ATCC 17978 with pAB3 plasmid S/T resistant NCBI accession number CP012004.1 | (1) |
| UPAB1 | Uropathogenic *A. baumannii* clinical isolate | (2) |
| AB347 | *A. baumannii*, respiratory clinical isolate from Bolivia (2016) | (3) |
| AB431 | *A. baumannii*, urinary clinical isolate from United States (2019) | Atlanta |
| AB774 | *A. baumannii*, respiratory clinical isolate from France (2022) | France |
| ARC6851 | *A. baumannii* | (4) |
| ARC6851 Δ*pilA* | ARC6851 *pilA* mutant | This study |
| AR0129 | *K. pneumoniae* | CDC Panel Name Enterobacterales Carbapenemase Diversity (CRE) AR Bank Number 0129 |
| AR0046 | *K. pneumoniae* | CDC Panel Gram Negative Carbapenemase Detection (CarbaNP) AR Bank Number 0046 |
| AR0126 | *K. pneumoniae* | CDC Panel Name Enterobacterales Carbapenemase Diversity (CRE) AR Bank Number 0126 |
| KR49 | *K. pneumoniae* | CDC KR Bank Number 49 |
| AR0125 | *K. pneumoniae* | CDC Panel Name Enterobacterales Carbapenemase Diversity (CRE) AR Bank Number 0125 |
| AR0073 | *E. cloacae* | CDC Panel Gram Negative Carbapenemase Detection (CarbaNP) AR Bank Number 0073 |
| 409957 | *P. aeruginosa* | (5) |
| 369569 | *P. aeruginosa* | (5) |
| 358800 | *P. aeruginosa* | (5) |
| *S. aureus* | *S. aureus* | Newman |

**Table S2: Plasmids used in this study.**

[a]Apr, apramycin; Zeo, zeocin

| **Plasmid** | **Description**[a] | **Source** |
|---|---|---|
| pKD4-Apr | Source for Apramycin cassette for mutant generation, $Apr^r$ | (4) |
| pAT03 | pMMB67EH with FLP recombinase | (6) |
| pAT04 | pMMB67EH with RecAbsystem, Hyg | (6) |
| pUCT18T-miniTn7T-Zeo | mTn7 complementation vector, $Zeo^r$ | (7) |
| pUCT18T-miniTn7T-Zeo-ARC_*pilA* | ARC6851 Δ*pilA* complementation construct, $Zeo^r$ | This study |

1. Weber BS, Ly PM, Irwin JN, Pukatzki S, Feldman MF. 2015. A multidrug resistance plasmid contains the molecular switch for type VI secretion in Acinetobacter baumannii. Proc Natl Acad Sci U S A 112:9442-7.
2. Di Venanzio G F-mA, Calix JJ, Haurat MF, Scott NE, Palmer LD, Potter RF, Hibbing ME, Friedman L, Wang B, Dantas G, Skaar EP, Hultgren SJ, Feldman MF. 2019. Urinary tract colonization is enhanced by a plasmid that regulates uropathogenic Acinetobacter baumannii chromosomal genes. Nat Commun 10:1–13.
3. Cerezales M, Xanthopoulou K, Wille J, Bustamante Z, Seifert H, Gallego L, Higgins PG. 2019. Acinetobacter baumannii analysis by core genome multi-locus sequence typing in two hospitals in Bolivia: endemicity of international clone 7 isolates (CC25). Int J Antimicrob Agents 53:844-849.
4. McGuffey JC, Jackson-Litteken CD, Di Venanzio G, Zimmer AA, Lewis JM, Distel JS, Kim KQ, Zaher HS, Alfonzo J, Scott NE, Feldman MF. 2023. The tRNA methyltransferase TrmB is critical for Acinetobacter baumannii stress responses and pulmonary infection. mBio doi:10.1128/mbio.01416-23:e0141623.
5. Lebreton F, Snesrud E, Hall L, Mills E, Galac M, Stam J, Ong A, Maybank R, Kwak YI, Johnson S, Julius M, Ly M, Swierczewski B, Waterman PE, Hinkle M, Jones A, Lesho E, Bennett JW, McGann P. 2021. A panel of diverse Pseudomonas aeruginosa clinical isolates for research and development. JAC Antimicrob Resist 3:dlab179.
6. Tucker AT, Nowicki EM, Boll JM, Knauf GA, Burdis NC, Trent MS, Davies BW. 2014. Defining gene-phenotype relationships in Acinetobacter baumannii through one-step chromosomal gene inactivation. mBio 5:e01313-14.
7. Ducas-Mowchun K, De Silva PM, Crisostomo L, Fernando DM, Chao TC, Pelka P, Schweizer HP, Kumar A. 2019. Next Generation of Tn7-Based Single-Copy Insertion Elements for Use in Multi- and Pan-Drug-Resistant Strains of Acinetobacter baumannii. Appl Environ Microbiol 85.

**Table S3: Primers used in this study**

| Primers | Sequence |
|---|---|
| F_U_*pilA*_1000bp_KO | TGCTGGCGTAGTAATGAG |
| R_U_*pilA*_1000bp_KO | CCAGCCTACACAATCGCTATTCATAGCCTTTTCCCC |
| F_D_*pilA*_1000bp_KO | AAGGAGGATATTCATATGTTCTGCTTGCCCTGC |
| R_D_*pilA*_1000bp_KO | CCACATCCCTATACGC |
| F_prom(500bp)_pilA_puCT18T-Zeo | GAGAAGCTTGGGCCCGGTACCGTAAGTCGATTGTAGAGCAGC |
| R_prom(500bp)_pilA_puCT18T-Zeo | GCAAGGCCTTCGCGAGGTACCTTATGCTGCAGGGCAAC |
| F_*pilA*_ARC6851_qRT-PCR | ACTCATGATCGTAGTTGCCATT |
| R_*pilA*_ARC6851_qRT-PCR | TTCACTAACCGCACGTGATAC |
| F_*pilA*_347_qRT-PCR | TGGTTGCCATTATCGGTATCT T |
| R_*pilA*_347_qRT-PCR | GGACAGCTGCTGCTACATTA |
| F_*rpoB*_qRT-PCR | ACGGTACTGAGCGTGTAATC |
| R_*rpoB*_qRT-PCR | TTACCACTTGAGTGGGTCTTAC |

| | MIC (µg/ml) | |
|---|---|---|
| Strain | DMSO | DF |
| *A. baumannii* ATCC 17978 | 2 | 1 |
| *A. baumannii* UPAB1 | 4 or 8 | 2 |
| *A. baumannii* AB347 | 64 | 16 |
| *A. baumannii* AB431 | 64 | 16 |
| *A. baumannii* AB774 | 128 | 32 |
| *A. baumannii* ARC6851 | >1024 | 256 |
| *K. pneumoniae* AR0129 | 16 or 32 | 4 |
| *K. pneumoniae* AR0046 | 32 | 4 |
| *K. pneumoniae* AR0126 | 32 | 4 |
| *K. pneumoniae* KR49 | 128 | 32 |
| *K. pneumoniae* AR0125 | 256 | 32 |
| *E. cloacae* AR0073 | 16 | 4 |
| *P. aeruginosa* 409957 | 64 | 8 |
| *P. aeruginosa* 369569 | 32 | 8 |
| *P. aeruginosa* 358800 | 128 | 64 |
| *S. aureus* | >1024 | >1024 |

**Table S4: Diclofenac affects colistin susceptibility of Gram-negative pathogens.** Gram-negative and Gram-positive bacteria were screened for changes in MICs to colistin in combination with the solvent control DMSO or 100 µM diclofenac using a 2-fold broth dilution method. MIC was determined as <10% growth compared to a non-treated culture. DF (diclofenac).

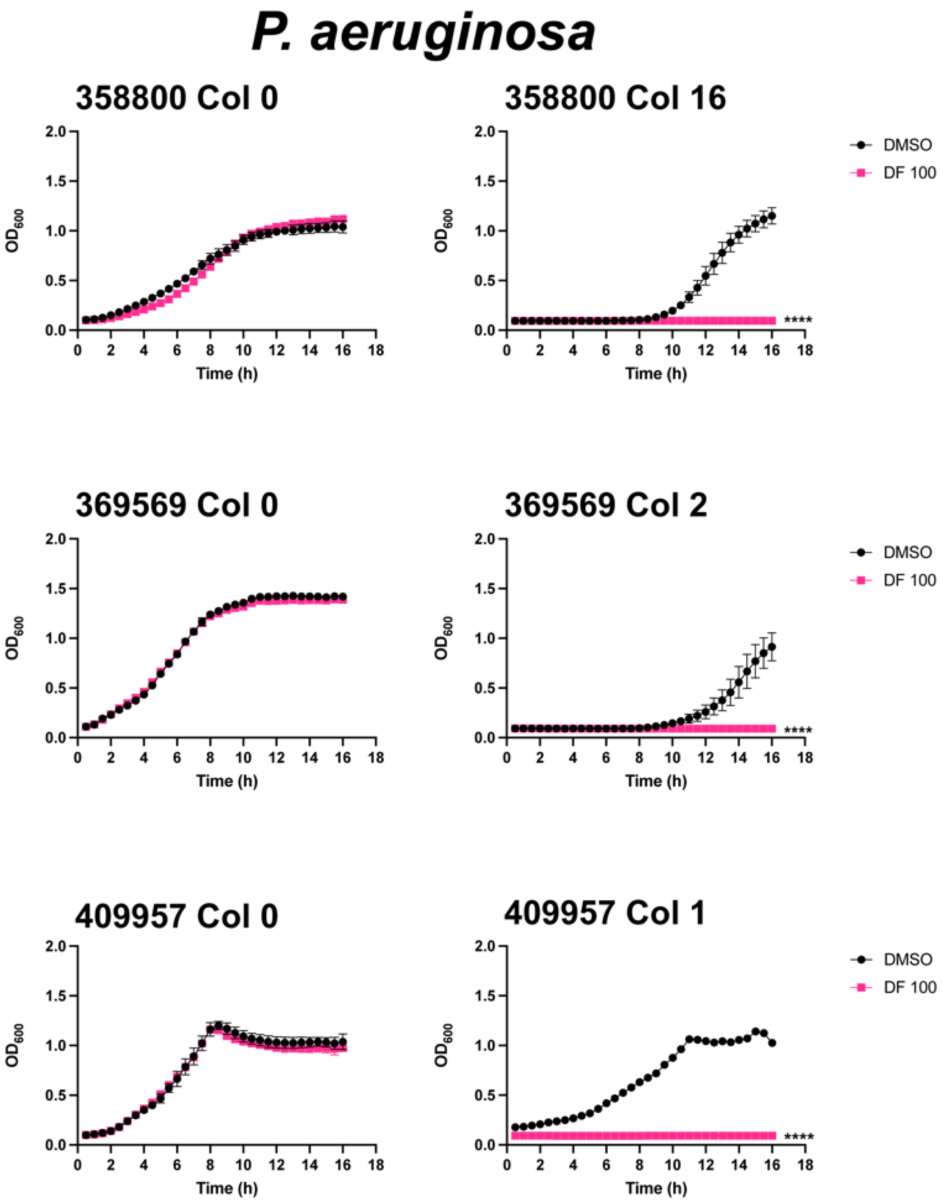


**Figure S1: Diclofenac inhibits *P. aeruginosa* growth in the presence of colistin.** Representative growth curves of *P. aeruginosa* 358800, 369569, and 409957 strains in LB containing either the solvent control DMSO or 100 µM diclofenac (DF) (left panel) and LB containing 16 µg/ml, 2 µg/ml, or 1 µg/ml colistin respectively in the presence of DMSO or 100 µM diclofenac (right panel). ****$P<0.0001$, unpaired *t* tests at 16 h for Col + DF 100 compared to DMSO control. Col (colistin), DF (diclofenac).

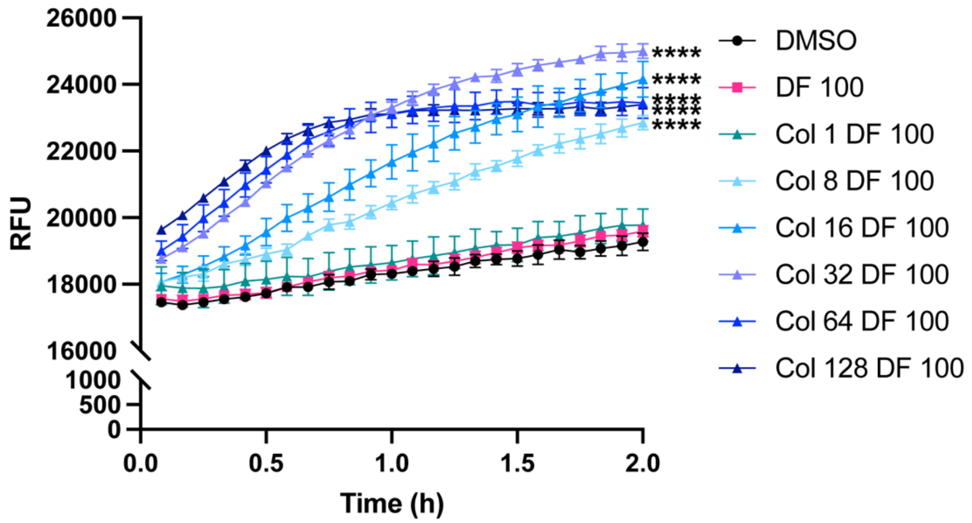


**Figure S2: Colistin, but not diclofenac, increases membrane permeability of ARC6851.** Representative membrane permeability assay measuring uptake of Hoescht 33342 (H33342) fluorescent dye. Bacterial cells were suspended in PBS supplemented with increasing concentrations of colistin (Col 1 µg/ml, 8 µg/ml, 16 µg/ml, 32 µg/ml, 64 µg/ml, or 128 µg/ml) in combination with the solvent control DMSO or 100 µM DF. Relative fluorescent units (RFU) were monitored over the course of 2 hours. ****$P$<0.0001, ns=not significant (One-way ANOVA at 16h with Tukey's test for multiple comparisons, results were compared to the control group treated with DMSO). Col (colistin), DF (diclofenac).

| | ARC6851 | | |
|---|---|---|---|
| **Treatment** | **DMSO** | **Oleic Acid 20 µg/ml** | **Linoleic Acid 20 µg/ml** |
| **Triclosan 0.05 µg/ml** | **32** | **512** | **256** |

**Table S5: Addition of exogenous oleic acid and linoleic acid abolishes synergy between colistin and triclosan.** ARC6851 was screened for changes in MICs to triclosan in combination with either the solvent control DMSO or 20 µg/ml of oleic acid and linoleic acid, using a 2-fold broth dilution method. MIC was determined as <10% growth compared to a non-treated culture.

| | ARC6851 | |
|---|---|---|
| **Treatment** | **Col Fatty Acid** | **Col DF Fatty Acid** |
| **Palmitic Acid** | **>1024** | **256** |
| **Araquidonic Acid** | **>1024** | **256** |
| **Linoleic Acid** | **>1024** | **256** |
| **Oleic Acid** | **>1024** | **512** |
| **Stearic Acid** | **>1024** | **512** |

**Table S6: Addition of exogenous fatty acids does not abolish synergy between colistin and diclofenac.** ARC6851 was screened for changes in MICs to colistin in combination with either the solvent control DMSO or 20 μg/ml of palmitic acid, araquidonic acid, linoleic acid, oleic acid, and stearic acid using a 2-fold broth dilution method. MIC was determined as <10% growth compared to a non-treated culture. Col (colistin), DF (diclofenac).

| | ARC6851 | | UPAB1 | |
|---|---|---|---|---|
| **Antibiotic** | **DMSO** | **DF** | **DMSO** | **DF** |
| Ciprofloxacin | 128 | 64 | 64 | 64 |
| Erythromycin | 32 or 64 | 32 or 64 | 32 | 32 |
| Tetracycline | 8 | 8 | 512 | 512 |
| Vancomycin | 128 or 256 | 128 or 256 | 512 | 512 |
| Imipenem | 128 or 256 | 128 or 256 | 256 | 256 |
| Sulfomethoxazole/ Trimethoprim | 960/192 | 960/192 | 480/96 | 480/96 |

**Table S7: Diclofenac does not affect *A. baumannii* susceptibility to ciprofloxacin, erythromycin, tetracycline, vancomycin, imipenem or sulfomethoxazole/trimethoprim.** ARC6851 and UPAB1 were screened for changes in MICs to ciprofloxacin, erythromycin, tetracycline, vancomycin, imipenem, or sulfomethoxazole/trimethoprim in combination with the solvent control DMSO or 100 μM diclofenac using a 2-fold broth dilution method. MIC was determined as <10% growth compared to a non-treated culture. DF (diclofenac).

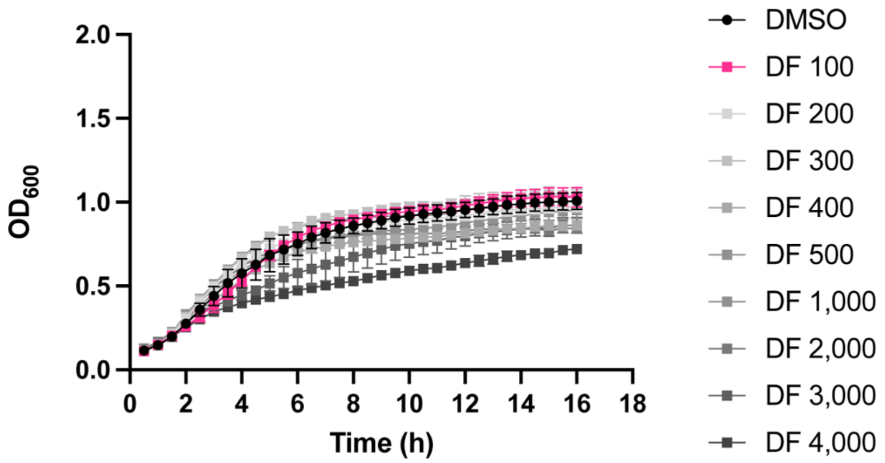


**Figure S3**: **Diclofenac does not effect ARC6851 growth at concentrations up to 4,000 µM.** Representative growth curves of ARC6851 in LB containing increasing concentrations of diclofenac (100 µM, 200 µM, 300 µM, 400 µM, 500 µM, 1,000 µM, 2,000 µM, 3,000 µM, or 4000 µM). DF (diclofenac).

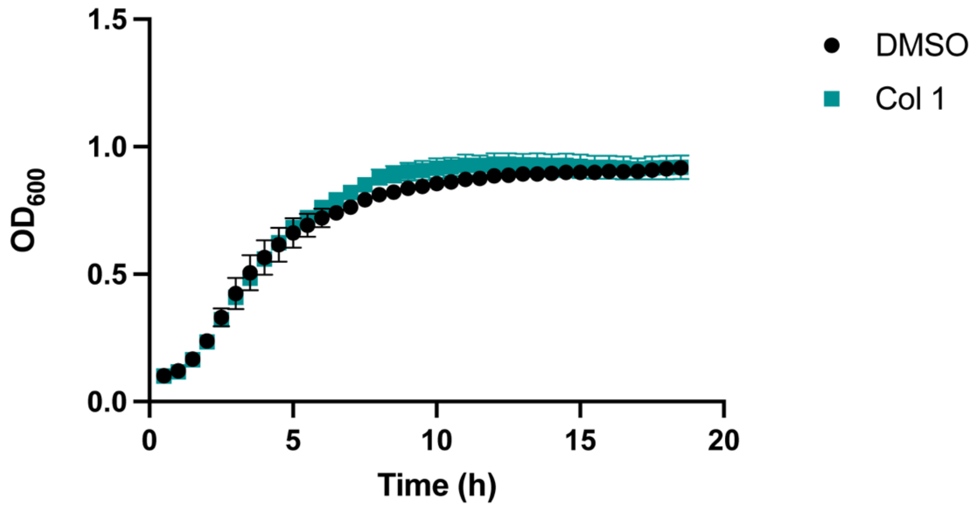


**Figure S4**: **Colistin at 1 µg/ml does not affect ARC6851 growth.** Representative growth curves of ARC6851 in LB containing either the solvent control DMSO or 1 µg/ml colistin. Col (colistin)

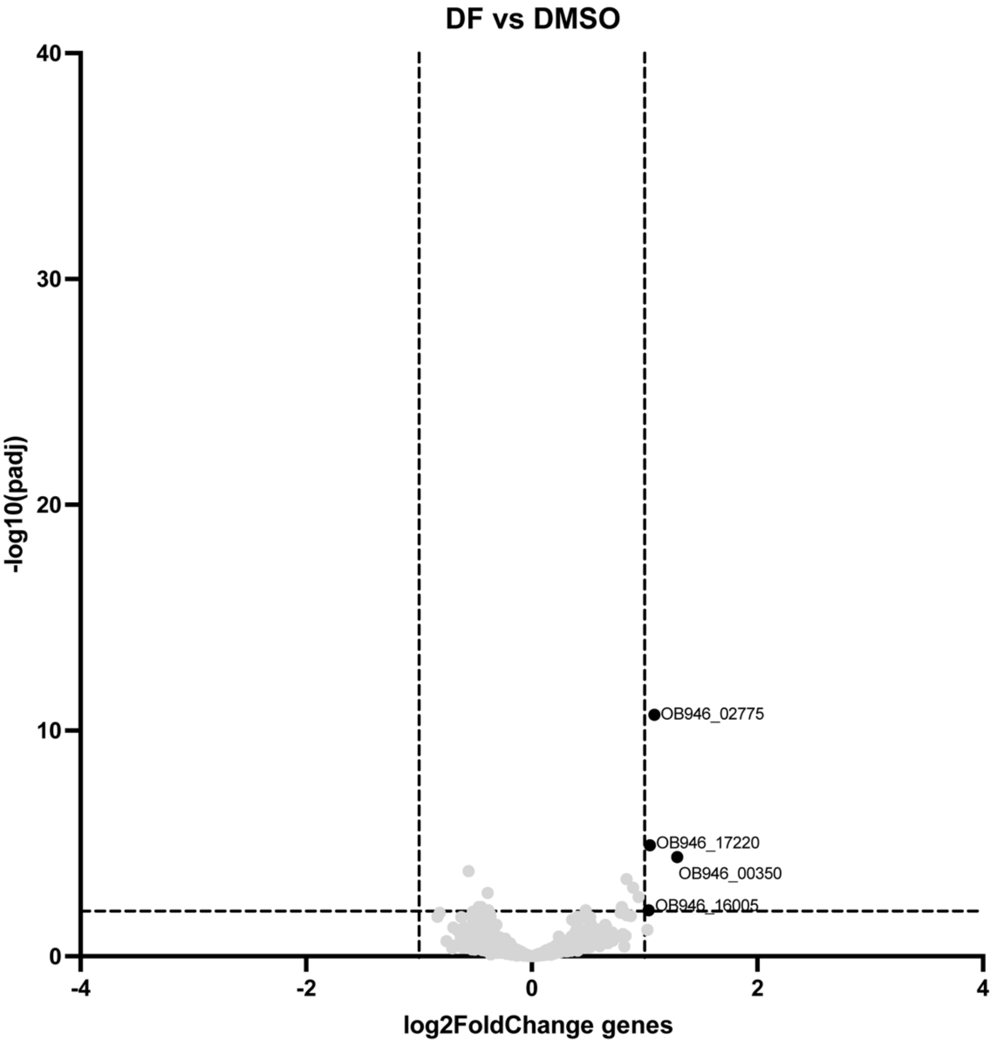


**Figure S5: Lipid metabolism and homeostasis genes upregulated under diclofenac treatment alone.** Volcano plot of Supplemental data set showing differentially expressed genes in ARC6851 in diclofenac treatment vs DMSO control. DF (diclofenac).

**Table S8: Upregulated genes in ARC6851 in diclofenac treatment vs DMSO.**

| Accession | Annotated gene | Fold change[a] |
|---|---|---|
| OB946_00350 | fahA | 2.44 |
| OB946_02775 | Acyl-CoA dehydrogenase C-terminal domain-containing protein | 2.12 |
| OB946_17220 | fadB | 2.06 |
| OB946_16005 | Lipid transport and metabolism | 2.05 |

a| Fold change cutoff: 2-fold *P* value < 0.01. Differential expression was calculated with DESeq2.

# A

**Genes differentially regulated**

| Condition | Number of genes |
|---|---|
| Colistin | 0 |
| Diclofenac | 4 |
| Colistin + Diclofenac | 130 |

# B

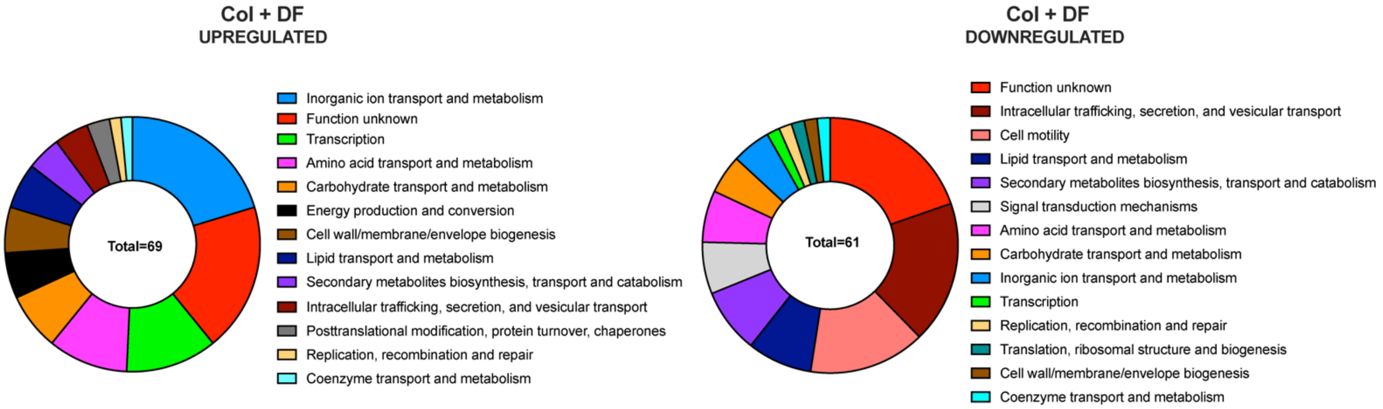


**Figure S6: Functional analysis of colistin and diclofenac treatment in ARC6851. (A)** Number of differentially regulated genes in ARC6851 when treated with 1μg/ml colistin, 100 μM diclofenac, or in combination. **(B)** Donut charts representing the functional classifications of differentially regulated genes of the combination treatment against DMSO. Classifications were determined based on the designated clusters of orthologous groups (COGs) using eggNOG Mapper. Col (colistin), DF (diclofenac).

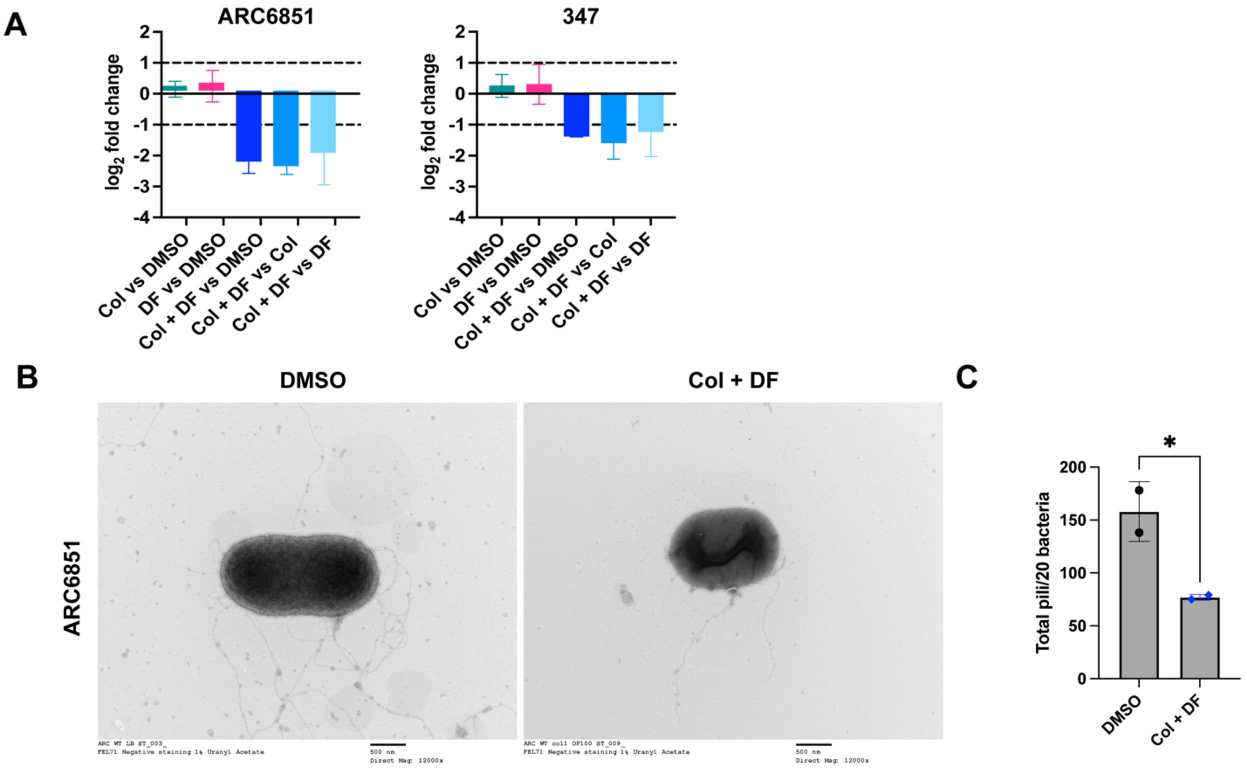


**Figure S7: Type IV pili decreases in ARC6851 under colistin and diclofenac combination treatment. (A)** Relative gene expression of *pilA* in ARC6851 grown in LB + DMSO, LB+ colistin (1 µg/ml) + DMSO, LB + diclofenac (100 µM), or LB + colistin (1 µg/ml) + diclofenac (100 µM) as determined by qRT-PCR. Dotted lines represent 2-fold change. **(B)** Transmission electron microscopy of ARC6851 WT in LB plus DMSO and LB plus colistin (1 µg/ml) plus diclofenac (100 µM). Scale bar of 500 nm is shown. **(C)** Quantification of type IV pili in ARC6851 WT was performed in 20 bacterial cells. Col (colistin), DF (diclofenac). Unpaired *t* tests *$P < 0.05$.

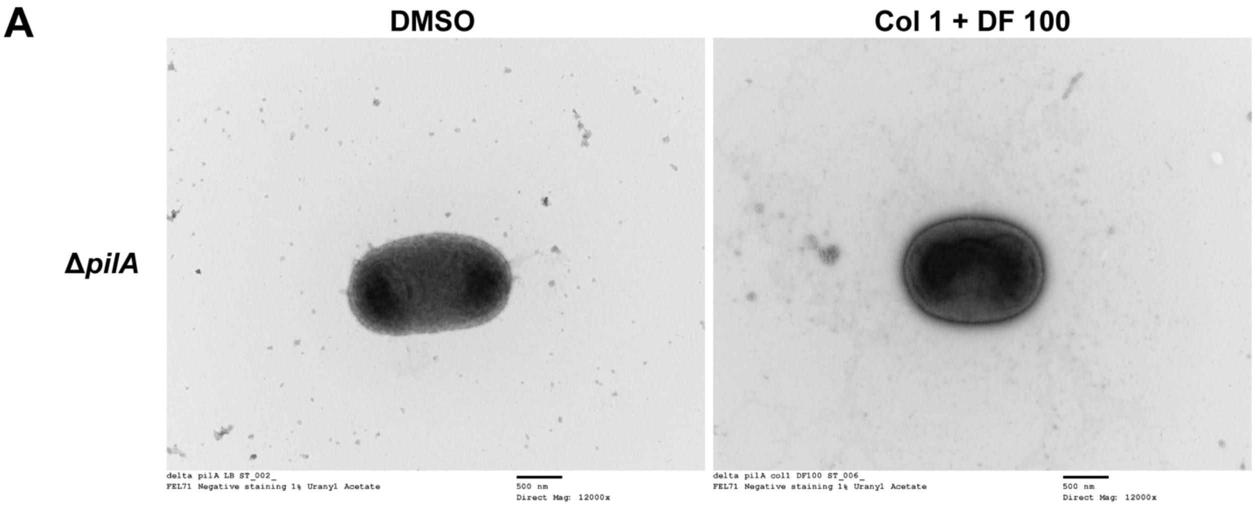


**B**

| | MIC (µg/ml) | |
|---|---|---|
| **Strain** | **DMSO** | **DF** |
| ***A. baumannii*** **ARC6851 Δ*pilA*** | **>1024** | **512** |

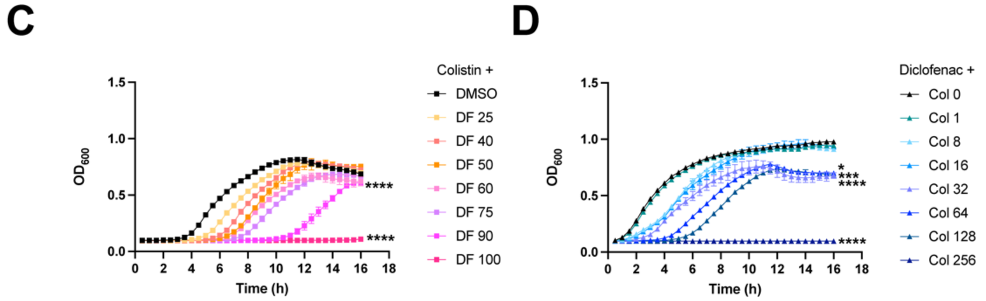


**Figure S8: Colistin and diclofenac kill ARC6851 Δ*pilA* in a synergistic manner. (A)** Transmission electron microscopy of ARC6851 Δ*pilA* in LB plus DMSO and LB plus colistin (1 µg/ml) plus diclofenac (100 µM). Scale bar of 500 nm is shown. **(B)** Δ*pilA* was screened for changes in MICs to colistin in combination with the solvent control DMSO and 100 µM diclofenac using a 2-fold broth dilution method. MIC was determined as <10% growth compared to a non-treated culture. **(C)** Representative growth curves of ARC6851 Δ*pilA* in LB containing 256 µg/ml colistin with increasing concentrations of diclofenac (25 µM, 40 µM, 50 µM, 60 µM, 75 µM, 90 µM, or 100 µM). **(D)** Representative growth curves of ARC6851 Δ*pilA* in LB containing 100 µM diclofenac with increasing concentrations of colistin (1 µg/ml, 8 µg/ml, 16 µg/ml, 32 µg/ml, 64 µg/ml, 128 µg/ml, or 256 µg/ml). *$P$<0.05, ***$P$<0.001, ****$P$<0.0001

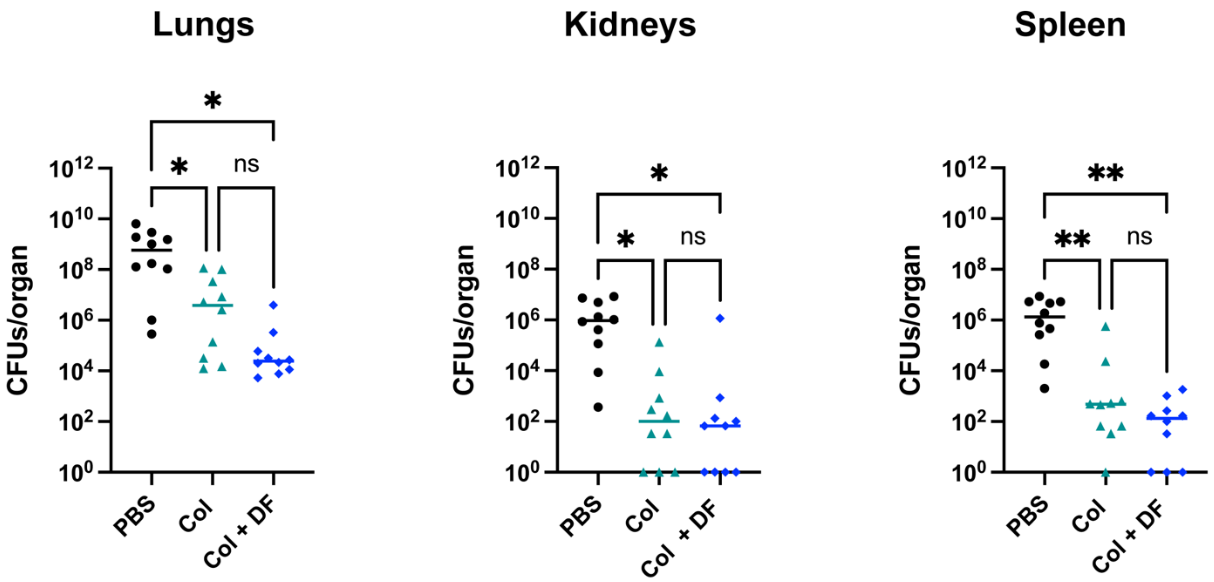


**Figure S9: ARC6851 Δ*pilA* is not attenuated in an acute murine pneumonia model.** C57BL/6 mice were infected with ~5 × $10^7$ CFU of mid-exponential ARC6851 WT or Δ*pilA*. At 24 h post-infection, the lungs, kidneys, and spleens were harvested, and the bacterial load present in each tissue was determined with serial dilutions. Each symbol represents an individual mouse, and the horizontal bar represents the median. Data collected from two independent experiments. *$P < 0.05$, **$P < 0.01$, Kruskal-Wallis test.

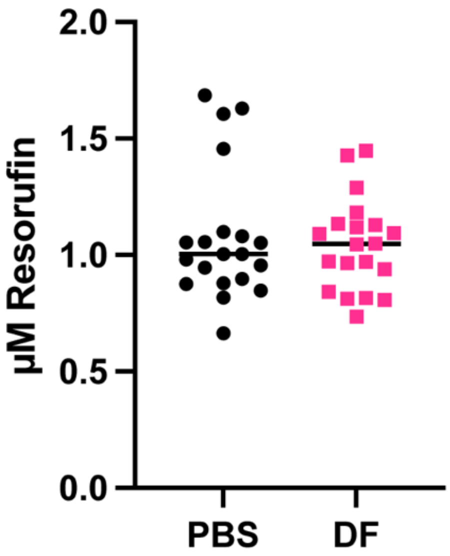


**Figure S10: Cycloxygenase COX-2 activity is not increased under diclofenac treatment.** Lung lysates containing protease inhibitors were used to determine COX-2 activities using a COX fluorescent activity assay kit. Resorufin fluorescence can be analyzed with an excitation wavelength between 530-540 nm and an emission wavelength between 585-595 nm. DF (diclofenac).

**Supplemental data set 1: Differentially expressed genes in ARC6851 in colistin + diclofenac treatment vs DMSO.**

| Accession | Annotated gene | Fold change[a] |
|---|---|---|
| **OB946_12785** | **NAD(P)/FAD-dependent oxidoreductase** | **11.37** |
| **OB946_12790** | **SDR family oxidoreductase** | **10.61** |
| **OB946_00885** | **FMN reductase** | **6.11** |
| **OB946_18865** | **RcnB family protein** | **5.64** |
| OB946_04015 | TetR/AcrR family transcriptional regulator | 5.56 |
| OB946_01760 | hypothetical protein | 5.52 |
| OB946_09560 | hypothetical protein | 4.99 |
| OB946_15220 | TetR/AcrR family transcriptional regulator | 4.68 |
| **OB946_02760** | **RcnB family protein** | **4.65** |
| OB946_12795 | alpha/beta hydrolase | 4.29 |
| OB946_02780 | hypothetical protein | 4.11 |
| **OB946_15490** | **MFS transporter** | **4.00** |
| **OB946_10710** | **SfnB family sulfur acquisition oxidoreductase** | **3.65** |
| OB946_19645 | | 3.50 |
| OB946_09895 | sulfite exporter TauE/SafE family protein | 3.48 |
| **OB946_18890** | **aliphatic sulfonate ABC transporter permease SsuC** | **3.31** |
| **OB946_00880** | **dimethyl sulfone monooxygenase SfnG** | **3.18** |
| **OB946_18885** | **FMNH2-dependent alkanesulfonate monooxygenase** | **3.11** |
| OB946_09155 | hypothetical protein | 3.11 |
| **OB946_18880** | **sulfonate ABC transporter substrate-binding protein** | **3.11** |
| OB946_08735 | MBL fold metallo-hydrolase | 3.01 |
| **OB946_02885** | **peroxiredoxin** | **2.91** |
| **OB946_10965** | **monooxygenase** | **2.88** |
| **OB946_18860** | **RcnB family protein** | **2.87** |
| OB946_09715 | lipoyl synthase | 2.81 |
| OB946_10655 | hypothetical protein | 2.79 |
| OB946_17300 | hypothetical protein | 2.78 |
| OB946_08740 | TetR family transcriptional regulator | 2.77 |
| OB946_18980 | hypothetical protein | 2.73 |
| **OB946_16170** | **MacA family efflux pump subunit** | **2.70** |
| **OB946_09580** | **MFS transporter** | **2.67** |
| OB946_00905 | DUF485 domain-containing protein | 2.61 |
| OB946_17670 | signal peptidase II | 2.55 |
| OB946_15495 | substrate-binding domain-containing protein | 2.54 |
| OB946_07115 | hypothetical protein | 2.54 |
| OB946_01830 | TIGR03915 family putative DNA repair protein | 2.50 |
| **OB946_10945** | **taurine ABC transporter substrate-binding protein** | **2.50** |
| OB946_18985 | hypothetical protein | 2.47 |
| **OB946_10720** | **LLM class flavin-dependent oxidoreductase** | **2.44** |
| OB946_18255 | hypothetical protein | 2.42 |
| OB946_19015 | M57 family metalloprotease | 2.42 |
| OB946_10755 | Crp/Fnr family transcriptional regulator | 2.40 |
| **OB946_18240** | **MFS transporter** | **2.39** |
| **OB946_03935** | **multidrug efflux RND transporter permease subunit** | **2.39** |
| OB946_14280 | phage tail protein | 2.39 |

| | | |
|---|---|---|
| OB946_10925 | aspartate/glutamate racemase family protein | 2.39 |
| OB946_18645 | GntR family transcriptional regulator | 2.38 |
| **OB946_13205** | **cation diffusion facilitator family transporter** | **2.34** |
| OB946_02765 | phosphate-starvation-inducible PsiE family protein | 2.34 |
| OB946_13200 | LysE/ArgO family amino acid transporter | 2.32 |
| **OB946_15225** | **MFS transporter** | **2.22** |
| OB946_19640 | | 2.21 |
| OB946_14925 | lipoprotein insertase outer membrane protein LolB | 2.20 |
| OB946_09705 | alpha-ketoacid dehydrogenase subunit beta | 2.20 |
| OB946_15450 | DUF1852 domain-containing protein | 2.17 |
| **OB946_09500** | **MFS transporter** | **2.16** |
| OB946_02270 | DUF3108 domain-containing protein | 2.16 |
| OB946_17295 | YMGG-like glycine zipper-containing protein | 2.15 |
| **OB946_04415** | **sulfate ABC transporter permease subunit CysT** | **2.13** |
| OB946_13800 | BCCT family transporter | 2.13 |
| OB946_18370 | PadR family transcriptional regulator | 2.13 |
| **OB946_10940** | **ATP-binding cassette domain-containing protein** | **2.12** |
| **OB946_04885** | **phosphate ABC transporter permease subunit PstC** | **2.11** |
| OB946_01140 | hypothetical protein | 2.11 |
| OB946_16770 | LysR family transcriptional regulator | 2.11 |
| OB946_16295 | class I SAM-dependent methyltransferase | 2.10 |
| **OB946_09460** | **anthranilate 1,2-dioxygenase small subunit** | **2.09** |
| OB946_19635 | DUF3861 domain-containing protein | 2.08 |
| OB946_18365 | DUF3861 domain-containing protein | 2.08 |
| OB946_10995 | LysR family transcriptional regulator | 2.08 |
| **OB946_00330** | **4-hydroxyphenylpyruvate dioxygenase** | **2.05** |
| OB946_03965 | transglycosylase SLT domain-containing protein | 2.05 |
| **OB946_04410** | **sulfate ABC transporter permease subunit CysW** | **2.05** |
| OB946_18195 | hypothetical protein | 2.04 |
| OB946_08485 | DUF6438 domain-containing protein | 0.49 |
| OB946_17110 | type II secretion system F family protein | 0.49 |
| OB946_15850 | hypothetical protein | 0.48 |
| OB946_01575 | VWA domain-containing protein | 0.48 |
| OB946_08640 | catechol 1,2-dioxygenase | 0.48 |
| OB946_04395 | ornithine uptake porin CarO type 1 | 0.48 |
| OB946_08695 | alpha/beta hydrolase | 0.47 |
| OB946_17055 | Na+/H+ antiporter subunit G | 0.47 |
| OB946_04995 | EAL domain-containing protein | 0.46 |
| OB946_09245 | hypothetical protein | 0.46 |
| OB946_08230 | cyd operon YbgE family protein | 0.46 |
| OB946_09660 | FAD/NAD(P)-binding oxidoreductase | 0.46 |
| OB946_09670 | MBL fold metallo-hydrolase | 0.46 |
| **OB946_01455** | **type IV pilus secretin PilQ family protein** | 0.45 |
| OB946_03135 | methyl-accepting chemotaxis protein | 0.44 |
| **OB946_11650** | **Paal family thioesterase** | **0.44** |
| OB946_11195 | LysE family translocator | 0.44 |
| OB946_08645 | muconolactone Delta-isomerase | 0.43 |

| | | |
|---|---|---|
| OB946_07615 | benzoate/H(+) symporter BenE family transporter | 0.43 |
| OB946_19040 | DMT family protein | 0.43 |
| **OB946_10590** | **fimbrial protein** | 0.43 |
| OB946_05255 | DUF3015 family protein | 0.43 |
| OB946_15250 | mechanosensitive ion channel | 0.42 |
| OB946_03145 | hypothetical protein | 0.41 |
| OB946_18220 | DNA-processing protein DprA | 0.41 |
| **OB946_01560** | **type IV pilus modification protein PilV** | 0.41 |
| OB946_04335 | multidrug effflux MFS transporter | 0.41 |
| OB946_12235 | DUF4882 family protein | 0.40 |
| OB946_04785 | type I-F CRISPR-associated endoribonuclease Cas6/Csy4 | 0.40 |
| **OB946_11660** | **PaaX** | **0.40** |
| **OB946_01565** | **PilW family protein** | 0.38 |
| OB946_15885 | 50S ribosomal protein L35 | 0.38 |
| OB946_10625 | hypothetical protein | 0.37 |
| **OB946_01450** | **pilus assembly protein PilP** | 0.36 |
| **OB946_11685** | **enoyl-CoA hydratase-related protein** | **0.36** |
| OB946_09310 | hypothetical protein | 0.36 |
| OB946_00200 | MFS transporter | 0.36 |
| OB946_04780 | type I-F CRISPR-associated protein Csy3 | 0.35 |
| OB946_08785 | hydrolase | 0.35 |
| **OB946_01555** | **GspH/FimT family pseudopilin** | 0.35 |
| **OB946_11680** | **2-(1,2-epoxy-1,2-dihydrophenyl)acetyl-CoA** | **0.35** |
| OB946_03140 | isomerase PaaG | 0.34 |
| **OB946_11665** | **phenylacetate--CoA ligase PaaK** | **0.34** |
| OB946_13975 | hypothetical protein | 0.33 |
| OB946_01570 | hypothetical protein | 0.33 |
| OB946_03165 | entericidin A/B family lipoprotein | 0.32 |
| OB946_09665 | protein tyrosine phosphatase family protein | 0.30 |
| OB946_09285 | hypothetical protein | 0.28 |
| OB946_09225 | hypothetical protein | 0.28 |
| OB946_15000 | peptidoglycan-binding protein LysM | 0.27 |
| **OB946_11655** | **DapH/DapD/GlmU-related protein** | **0.26** |
| OB946_09280 | hypothetical protein | 0.26 |
| OB946_07585 | universal stress protein | 0.25 |
| **OB946_11675** | **3-hydroxyacyl-CoA dehydrogenase** | **0.25** |
| **OB946_14630** | **PilT/PilU family type 4a pilus ATPase** | 0.25 |
| OB946_10620 | OB946_10620 | 0.24 |
| **OB946_11670** | **3-oxoadipyl-CoA thiolase** | **0.19** |
| OB946_09205 | hypothetical protein | 0.19 |
| **OB946_01505** | **pilin** | 0.11 |
| OB946_09275 | major capsid protein | 0.08 |
| OB946_09200 | major capsid protein | 0.08 |

a| Fold change cutoff: 2-fold *P* value < 0.01. Differential expression was calculated with DESeq2.

**Supplemental data set 2: Differentially expressed genes in ARC6851 in colistin + diclofenac treatment vs colistin.**

| Accession | Annotated gene | Fold change[a] |
|---|---|---|
| **OB946_12790** | SDR family oxidoreductase | 11.90 |
| **OB946_15490** | MFS transporter | 10.24 |
| **OB946_12785** | NAD(P)/FAD-dependent oxidoreductase | 9.67 |
| **OB946_00885** | FMN reductase | 8.90 |
| **OB946_01760** | hypothetical protein | 7.10 |
| **OB946_18875** | sulfonate ABC transporter substrate-binding protein | 6.53 |
| **OB946_18475** | LuxR family transcriptional regulator AbaR | 6.03 |
| **OB946_19460** | | 5.84 |
| **OB946_12795** | alpha/beta hydrolase | 5.78 |
| **OB946_17865** | site-specific integrase | 5.50 |
| **OB946_06760** | hypothetical protein | 5.34 |
| **OB946_12665** | hypothetical protein | 5.30 |
| **OB946_15220** | TetR/AcrR family transcriptional regulator | 5.28 |
| **OB946_01260** | hypothetical protein | 5.24 |
| **OB946_18980** | hypothetical protein | 5.10 |
| **OB946_08890** | DMT family transporter | 4.96 |
| **OB946_12935** | hypothetical protein | 4.92 |
| **OB946_19570** | | 4.87 |
| **OB946_00905** | DUF485 domain-containing protein | 4.82 |
| **OB946_18885** | FMNH2-dependent alkanesulfonate monooxygenase | 4.73 |
| **OB946_08510** | LysR family transcriptional regulator | 4.61 |
| **OB946_09895** | sulfite exporter TauE/SafE family protein | 4.58 |
| **OB946_00715** | DUF1737 domain-containing protein | 4.57 |
| **OB946_17925** | phage virion morphogenesis protein | 4.30 |
| **OB946_18985** | hypothetical protein | 4.16 |
| **OB946_01140** | hypothetical protein | 4.05 |
| **OB946_01200** | DMT family transporter | 3.81 |
| **OB946_00880** | dimethyl sulfone monooxygenase SfnG | 3.80 |

| | | |
|---|---|---|
| **OB946_13255** | hypothetical protein | 3.74 |
| **OB946_02760** | RcnB family protein | 3.73 |
| **OB946_02885** | peroxiredoxin | 3.73 |
| **OB946_18865** | RcnB family protein | 3.70 |
| **OB946_10925** | aspartate/glutamate racemase family protein | 3.70 |
| **OB946_01155** | hypothetical protein | 3.70 |
| **OB946_08055** | Lrp/AsnC family transcriptional regulator GigD | 3.68 |
| **OB946_17775** | ogr/Delta-like zinc finger family protein | 3.67 |
| **OB946_11820** | ankyrin repeat domain-containing protein | 3.66 |
| **OB946_13190** | hypothetical protein | 3.65 |
| **OB946_15450** | DUF1852 domain-containing protein | 3.64 |
| **OB946_08435** | 3-carboxy-cis%2Ccis-muconate cycloisomerase | 3.61 |
| **OB946_16795** | acyl-CoA dehydrogenase family protein | 3.60 |
| **OB946_05125** | hypothetical protein | 3.56 |
| **OB946_14190** | hypothetical protein | 3.56 |
| **OB946_05795** | OmpW family outer membrane protein | 3.53 |
| **OB946_04425** | sulfate ABC transporter substrate-binding protein | 3.50 |
| **OB946_08865** | SLC13 family permease | 3.50 |
| **OB946_13205** | cation diffusion facilitator family transporter | 3.45 |
| **OB946_09560** | hypothetical protein | 3.41 |
| **OB946_16570** | winged helix-turn-helix transcriptional regulator | 3.39 |
| **OB946_08740** | TetR family transcriptional regulator | 3.38 |
| **OB946_11725** | APC family permease | 3.38 |
| **OB946_14340** | hypothetical protein | 3.35 |
| **OB946_18645** | GntR family transcriptional regulator | 3.31 |
| **OB946_19480** | | 3.30 |
| **OB946_05030** | LysR substrate-binding domain-containing protein | 3.30 |
| **OB946_12395** | cold-shock protein | 3.30 |
| **OB946_15495** | substrate-binding domain-containing protein | 3.13 |
| **OB946_17165** | DUF1656 domain-containing protein | 3.12 |
| **OB946_01485** | metal-dependent hydrolase | 3.10 |

| | | |
|---|---|---|
| **OB946_01170** | AraC family transcriptional regulator | 3.09 |
| **OB946_18370** | PadR family transcriptional regulator | 3.08 |
| **OB946_04015** | TetR/AcrR family transcriptional regulator | 3.06 |
| **OB946_00650** | 2-oxo-4-hydroxy-4-carboxy-5-ureidoimidazoline decarboxylase | 3.06 |
| **OB946_07120** | glycine zipper domain-containing protein | 3.06 |
| **OB946_17300** | hypothetical protein | 3.04 |
| **OB946_19015** | M57 family metalloprotease | 3.04 |
| **OB946_08465** | type I 3-dehydroquinate dehydratase | 3.02 |
| **OB946_14445** | hypothetical protein | 3.02 |
| **OB946_10945** | taurine ABC transporter substrate-binding protein | 3.00 |
| **OB946_14390** | hypothetical protein | 2.98 |
| **OB946_01700** | Holliday junction resolvase-like protein | 2.97 |
| **OB946_19680** | | 2.92 |
| **OB946_09580** | MFS transporter | 2.90 |
| **OB946_07115** | hypothetical protein | 2.89 |
| **OB946_18880** | sulfonate ABC transporter substrate-binding protein | 2.89 |
| **OB946_17470** | TetR/AcrR family transcriptional regulator | 2.87 |
| **OB946_15225** | MFS transporter | 2.86 |
| **OB946_12365** | minor capsid protein | 2.86 |
| **OB946_18890** | aliphatic sulfonate ABC transporter permease SsuC | 2.84 |
| **OB946_08735** | MBL fold metallo-hydrolase | 2.82 |
| **OB946_03440** | lysophospholipid acyltransferase family protein | 2.80 |
| **OB946_07605** | magnesium-translocating P-type ATPase | 2.79 |
| **OB946_13200** | LysE/ArgO family amino acid transporter | 2.78 |
| **OB946_17840** | hypothetical protein | 2.73 |
| **OB946_10425** | metalloregulator ArsR/SmtB family transcription factor | 2.73 |
| **OB946_02780** | hypothetical protein | 2.72 |
| **OB946_12500** | benzoate 1%2C2-dioxygenase electron transfer component BenC | 2.72 |
| **OB946_12240** | hypothetical protein | 2.70 |

| | | |
|---|---|---|
| **OB946_14925** | lipoprotein insertase outer membrane protein LolB | 2.69 |
| **OB946_10715** | SfnB family sulfur acquisition oxidoreductase | 2.69 |
| **OB946_14165** | hypothetical protein | 2.68 |
| **OB946_19640** | | 2.66 |
| **OB946_11430** | acyl-CoA dehydrogenase family protein | 2.65 |
| **OB946_00325** | DUF523 domain-containing protein | 2.64 |
| **OB946_08060** | kynureninase | 2.64 |
| **OB946_07760** | type II 3-dehydroquinate dehydratase | 2.63 |
| **OB946_03935** | multidrug efflux RND transporter permease subunit | 2.63 |
| **OB946_10635** | PACE efflux transporter | 2.62 |
| **OB946_10265** | SRPBCC family protein | 2.61 |
| **OB946_10920** | NCS1 family nucleobase:cation symporter-1 | 2.59 |
| **OB946_14385** | RusA family crossover junction endodeoxyribonuclease | 2.59 |
| **OB946_19605** | | 2.58 |
| **OB946_19305** | | 2.53 |
| **OB946_12330** | hypothetical protein | 2.52 |
| **OB946_19340** | | 2.52 |
| **OB946_17560** | PAS domain-containing sensor histidine kinase | 2.51 |
| **OB946_12390** | hypothetical protein | 2.51 |
| **OB946_16170** | MacA family efflux pump subunit | 2.50 |
| **OB946_14090** | PAAR domain-containing protein | 2.49 |
| **OB946_00625** | allantoicase | 2.49 |
| **OB946_19635** | | 2.48 |
| **OB946_12835** | 3-(3-hydroxy-phenyl)propionate transporter MhpT | 2.48 |
| **OB946_05565** | hypothetical protein | 2.47 |
| **OB946_00330** | 4-hydroxyphenylpyruvate dioxygenase | 2.47 |
| **OB946_16820** | fluoride efflux transporter CrcB | 2.46 |
| **OB946_03085** | DUF488 domain-containing protein | 2.46 |
| **OB946_06020** | DUF559 domain-containing protein | 2.46 |
| **OB946_06065** | putative metallopeptidase | 2.46 |
| **OB946_06145** | hypothetical protein | 2.45 |

| | | |
|---|---|---|
| **OB946_18860** | RcnB family protein | 2.45 |
| **OB946_03505** | hypothetical protein | 2.44 |
| **OB946_18255** | hypothetical protein | 2.43 |
| **OB946_08930** | LysR family transcriptional regulator | 2.42 |
| **OB946_05200** | acinetobactin biosynthesis bifunctional isochorismatase/aryl carrier protein BasF | 2.42 |
| **OB946_12275** | phage minor head protein | 2.42 |
| **OB946_00985** | AzlC family ABC transporter permease | 2.41 |
| **OB946_14655** | bacteriohemerythrin | 2.41 |
| **OB946_10645** | integrase family protein | 2.41 |
| **OB946_08150** | hypothetical protein | 2.41 |
| **OB946_07860** | hypothetical protein | 2.41 |
| **OB946_19065** | hypothetical protein | 2.41 |
| **OB946_01120** | helix-turn-helix domain-containing protein | 2.40 |
| **OB946_09530** | CoA transferase subunit A | 2.40 |
| **OB946_04025** | SDR family oxidoreductase | 2.40 |
| **OB946_03965** | transglycosylase SLT domain-containing protein | 2.39 |
| **OB946_16315** | gluconokinase | 2.39 |
| **OB946_11495** | carboxymuconolactone decarboxylase family protein | 2.39 |
| **OB946_17730** | | 2.38 |
| **OB946_02740** | DUF4184 family protein | 2.38 |
| **OB946_05080** | hypothetical protein | 2.38 |
| **OB946_01830** | TIGR03915 family putative DNA repair protein | 2.37 |
| **OB946_17675** | cation transporter | 2.37 |
| **OB946_19475** | | 2.37 |
| **OB946_16335** | acetate kinase | 2.36 |
| **OB946_08065** | amino acid permease | 2.36 |
| **OB946_04865** | Lrp/AsnC family transcriptional regulator | 2.36 |
| **OB946_03370** | fatty acid desaturase family protein | 2.36 |
| **OB946_00040** | DUF6091 family protein | 2.36 |
| **OB946_12440** | hypothetical protein | 2.35 |

| | | |
|---|---|---|
| **OB946_18240** | MFS transporter | 2.34 |
| **OB946_11915** | type VI secretion system contractile sheath small subunit | 2.33 |
| **OB946_19555** | | 2.33 |
| **OB946_10965** | monooxygenase | 2.32 |
| **OB946_11375** | acyl-CoA dehydrogenase family protein | 2.31 |
| **OB946_15410** | hypothetical protein | 2.30 |
| **OB946_16320** | gluconate:H+ symporter | 2.30 |
| **OB946_04410** | sulfate ABC transporter permease subunit CysW | 2.29 |
| **OB946_17930** | phage baseplate assembly protein V | 2.26 |
| **OB946_11230** | iron-containing redox enzyme family protein | 2.26 |
| **OB946_09715** | lipoyl synthase | 2.25 |
| **OB946_00825** | putative porin | 2.25 |
| **OB946_01125** | GNAT family N-acetyltransferase | 2.24 |
| **OB946_00910** | cation acetate symporter | 2.24 |
| **OB946_09705** | alpha-ketoacid dehydrogenase subunit beta | 2.24 |
| **OB946_11475** | 4-hydroxybenzoate 3-monooxygenase | 2.24 |
| **OB946_16800** | SfnB family sulfur acquisition oxidoreductase | 2.23 |
| **OB946_08145** | hypothetical protein | 2.21 |
| **OB946_15505** | FAD-dependent tricarballylate dehydrogenase TcuA | 2.21 |
| **OB946_01630** | hemerythrin domain-containing protein | 2.21 |
| **OB946_19390** | | 2.21 |
| **OB946_16175** | MacB family efflux pump subunit | 2.20 |
| **OB946_09150** | FAD-binding oxidoreductase | 2.20 |
| **OB946_09615** | fumarate reductase/succinate dehydrogenase flavoprotein subunit | 2.20 |
| **OB946_12355** | helix-turn-helix domain-containing protein | 2.20 |
| **OB946_13895** | LysR family transcriptional regulator | 2.19 |
| **OB946_08050** | GNAT family N-acetyltransferase | 2.19 |
| **OB946_09830** | hypothetical protein | 2.19 |
| **OB946_05365** | hypothetical protein | 2.19 |
| **OB946_01255** | acetyl-CoA hydrolase/transferase family protein | 2.18 |
| **OB946_12105** | Lrp/AsnC family transcriptional regulator | 2.18 |

| | | |
|---|---|---|
| **OB946_13915** | hypothetical protein | 2.17 |
| **OB946_11500** | NAD-dependent succinate-semialdehyde dehydrogenase | 2.17 |
| **OB946_11005** | malonate transporter subunit MadL | 2.16 |
| **OB946_19500** | | 2.16 |
| **OB946_06790** | paraquat-inducible protein A | 2.15 |
| **OB946_13430** | sulfate adenylyltransferase subunit CysD | 2.15 |
| **OB946_10640** | LysR family transcriptional regulator | 2.14 |
| **OB946_04840** | TetR/AcrR family transcriptional regulator | 2.14 |
| **OB946_03170** | DUF1853 family protein | 2.14 |
| **OB946_00940** | TetR/AcrR family transcriptional regulator | 2.13 |
| **OB946_00320** | MATE family efflux transporter | 2.13 |
| **OB946_01705** | hypothetical protein | 2.13 |
| **OB946_15470** | AraC family transcriptional regulator | 2.12 |
| **OB946_01615** | lipase secretion chaperone | 2.12 |
| **OB946_14805** | hypothetical protein | 2.12 |
| **OB946_11020** | biotin-independent malonate decarboxylase subunit gamma | 2.12 |
| **OB946_08590** | flavin reductase family protein | 2.12 |
| **OB946_18995** | LysR family transcriptional regulator | 2.11 |
| **OB946_00645** | hydroxyisourate hydrolase | 2.11 |
| **OB946_09500** | MFS transporter | 2.11 |
| **OB946_19655** | | 2.11 |
| **OB946_08015** | regulatory protein RecX | 2.11 |
| **OB946_11530** | flavin reductase family protein | 2.11 |
| **OB946_04655** | hypothetical protein | 2.11 |
| **OB946_02270** | DUF3108 domain-containing protein | 2.10 |
| **OB946_11775** | dihydrodipicolinate synthase family protein | 2.10 |
| **OB946_09455** | anthranilate 1%2C2-dioxygenase large subunit | 2.10 |
| **OB946_15230** | HlyD family secretion protein | 2.10 |
| **OB946_08085** | SulP family inorganic anion transporter | 2.10 |
| **OB946_12360** | MFS transporter | 2.10 |
| **OB946_04420** | alpha/beta hydrolase | 2.09 |

| | | |
|---|---|---|
| **OB946_08470** | right-handed parallel beta-helix repeat-containing protein | 2.09 |
| **OB946_19515** | | 2.08 |
| **OB946_12735** | NAD(P)/FAD-dependent oxidoreductase | 2.08 |
| **OB946_07435** | hypothetical protein | 2.08 |
| **OB946_07820** | MoaD/ThiS family protein | 2.08 |
| **OB946_18125** | esterase-like activity of phytase family protein | 2.08 |
| **OB946_00185** | hypothetical protein | 2.07 |
| **OB946_12810** | MFS transporter | 2.07 |
| **OB946_01740** | arginine N-succinyltransferase | 2.06 |
| **OB946_02925** | hypothetical protein | 2.06 |
| **OB946_17700** | | 2.05 |
| **OB946_11885** | hypothetical protein | 2.05 |
| **OB946_04580** | hypothetical protein | 2.05 |
| **OB946_19645** | | 2.05 |
| **OB946_13800** | BCCT family transporter | 2.05 |
| **OB946_16295** | class I SAM-dependent methyltransferase | 2.05 |
| **OB946_17295** | YMGG-like glycine zipper-containing protein | 2.05 |
| **OB946_13650** | MFS transporter | 2.05 |
| **OB946_10775** | L-valine transporter subunit YgaH | 2.04 |
| **OB946_07130** | type 1 glutamine amidotransferase domain-containing protein | 2.04 |
| **OB946_08045** | hypothetical protein | 2.04 |
| **OB946_04585** | hypothetical protein | 2.04 |
| **OB946_00430** | ATP-binding protein | 2.04 |
| **OB946_15235** | pyridoxal-phosphate dependent enzyme | 2.04 |
| **OB946_09315** | fimbrial protein | 2.03 |
| **OB946_09540** | TIGR00366 family protein | 2.03 |
| **OB946_14855** | LysR family transcriptional regulator | 2.03 |
| **OB946_11155** | amino acid ABC transporter permease | 2.02 |
| **OB946_08540** | MFS transporter | 2.02 |
| **OB946_16330** | phosphogluconate dehydratase | 2.02 |
| **OB946_06305** | multidrug efflux transcriptional repressor AdeL | 2.02 |

| | | |
|---|---|---|
| **OB946_11345** | PaaI family thioesterase | 2.01 |
| **OB946_11385** | SDR family oxidoreductase | 2.01 |
| **OB946_04240** | rhombotarget A | 2.01 |
| **OB946_02785** | sensor histidine kinase efflux regulator BaeS | 2.01 |
| **OB946_07565** | OmpW family outer membrane protein | 2.01 |
| **OB946_02970** | SMR family transporter | 2.01 |
| **OB946_06850** | | 2.00 |
| **OB946_13195** | LysR family transcriptional regulator ArgP | 2.00 |
| **OB946_12955** | Lrp/AsnC family transcriptional regulator | 2.00 |
| **OB946_13735** | nicotinamide riboside transporter PnuC | 2.00 |
| **OB946_14245** | hypothetical protein | -2.00 |
| **OB946_08240** | cytochrome d ubiquinol oxidase subunit II | -2.01 |
| **OB946_08245** | cytochrome ubiquinol oxidase subunit I | -2.01 |
| **OB946_08630** | 3-oxoacid CoA-transferase subunit B | -2.01 |
| **OB946_03775** | universal stress protein | -2.02 |
| **OB946_19130** | 50S ribosomal protein L34 | -2.02 |
| **OB946_02005** | 30S ribosomal protein S10 | -2.02 |
| **OB946_11905** | type VI secretion system tube protein Hcp | -2.03 |
| **OB946_11485** | NAD(P)H-dependent oxidoreductase | -2.03 |
| **OB946_16105** | hypothetical protein | -2.04 |
| **OB946_12560** | hypothetical protein | -2.04 |
| **OB946_19040** | DMT family protein | -2.04 |
| **OB946_03790** | carbamoyl-phosphate synthase large subunit | -2.04 |
| **OB946_15320** | NADH-quinone oxidoreductase subunit NuoH | -2.05 |
| **OB946_06180** | hypothetical protein | -2.05 |
| **OB946_08235** | cytochrome bd-I oxidase subunit CydX | -2.05 |
| **OB946_11955** | biotin carboxylase N-terminal domain-containing protein | -2.06 |
| **OB946_09945** | hypothetical protein | -2.06 |
| **OB946_11195** | LysE family translocator | -2.06 |
| **OB946_03095** | YegP family protein | -2.08 |
| **OB946_06320** | ABC transporter permease | -2.09 |

| | | |
|---|---|---|
| **OB946_05530** | hypothetical protein | -2.09 |
| **OB946_06925** | hypothetical protein | -2.10 |
| **OB946_15330** | NADH-quinone oxidoreductase subunit NuoF | -2.10 |
| **OB946_03020** | translational GTPase TypA | -2.10 |
| **OB946_17110** | type II secretion system F family protein | -2.10 |
| **OB946_15300** | NADH-quinone oxidoreductase subunit L | -2.10 |
| **OB946_15290** | NADH-quinone oxidoreductase subunit NuoN | -2.10 |
| **OB946_05505** | phage terminase large subunit | -2.12 |
| **OB946_04395** | ornithine uptake porin CarO type 1 | -2.12 |
| **OB946_05560** | hypothetical protein | -2.13 |
| **OB946_15325** | NADH-quinone oxidoreductase subunit NuoG | -2.13 |
| **OB946_01580** | type IV pilin protein | -2.14 |
| **OB946_04995** | EAL domain-containing protein | -2.14 |
| **OB946_09270** | virulence factor TspB C-terminal domain-related protein | -2.15 |
| **OB946_01455** | type IV pilus secretin PilQ family protein | -2.15 |
| **OB946_01850** | acyl-CoA dehydrogenase C-terminal domain-containing protein | -2.16 |
| **OB946_09225** | hypothetical protein | -2.17 |
| **OB946_05525** | Gp49 family protein | -2.17 |
| **OB946_05330** | hypothetical protein | -2.18 |
| **OB946_07930** | nitrogen regulation protein NR(I) | -2.18 |
| **OB946_01575** | VWA domain-containing protein | -2.19 |
| **OB946_10865** | chromate transporter | -2.19 |
| **OB946_18220** | DNA-processing protein DprA | -2.20 |
| **OB946_15850** | hypothetical protein | -2.20 |
| **OB946_01565** | PilW family protein | -2.20 |
| **OB946_02000** | type 1 glutamine amidotransferase | -2.20 |
| **OB946_01555** | GspH/FimT family pseudopilin | -2.21 |
| **OB946_09210** | hypothetical protein | -2.27 |
| **OB946_12535** | alkyl hydroperoxide reductase subunit C | -2.27 |
| **OB946_06675** | glycosyltransferase | -2.29 |
| **OB946_11225** | catalase HPII | -2.30 |

| | | |
|---|---|---|
| **OB946_03135** | methyl-accepting chemotaxis protein | -2.30 |
| **OB946_07655** | MaoC/PaaZ C-terminal domain-containing protein | -2.32 |
| **OB946_02010** | 50S ribosomal protein L3 | -2.32 |
| **OB946_09255** | hypothetical protein | -2.34 |
| **OB946_09285** | hypothetical protein | -2.35 |
| **OB946_08650** | muconate/chloromuconate family cycloisomerase | -2.36 |
| **OB946_13875** | Slam-dependent surface lipoprotein | -2.36 |
| **OB946_02020** | 50S ribosomal protein L23 | -2.37 |
| **OB946_07615** | benzoate/H(+) symporter BenE family transporter | -2.38 |
| **OB946_16150** | DUF4124 domain-containing protein | -2.40 |
| **OB946_13560** | hypothetical protein | -2.41 |
| **OB946_02015** | 50S ribosomal protein L4 | -2.42 |
| **OB946_09120** | multidrug efflux RND transporter periplasmic adaptor subunit AdeA | -2.43 |
| **OB946_15250** | mechanosensitive ion channel | -2.43 |
| **OB946_07650** | 3-oxoacyl-ACP reductase | -2.48 |
| **OB946_01450** | pilus assembly protein PilP | -2.50 |
| **OB946_11650** | PaaI family thioesterase | -2.50 |
| **OB946_14625** | type IV pilus twitching motility protein PilT | -2.51 |
| **OB946_08685** | aldehyde dehydrogenase (NADP(+)) | -2.53 |
| **OB946_04335** | multidrug effflux MFS transporter | -2.53 |
| **OB946_01050** | MFS transporter | -2.54 |
| **OB946_11660** | phenylacetic acid degradation operon negative regulatory protein PaaX | -2.54 |
| **OB946_11685** | enoyl-CoA hydratase-related protein | -2.54 |
| **OB946_15715** | ribosome-associated translation inhibitor RaiA | -2.55 |
| **OB946_10780** | tautomerase family protein | -2.56 |
| **OB946_05545** | hypothetical protein | -2.58 |
| **OB946_08230** | cyd operon YbgE family protein | -2.59 |
| **OB946_03145** | hypothetical protein | -2.63 |
| **OB946_08695** | alpha/beta hydrolase | -2.65 |
| **OB946_05540** | Ish1 domain-containing protein | -2.65 |
| **OB946_09165** | hypothetical protein | -2.73 |

| | | |
|---|---|---|
| **OB946_11680** | 2-(1%2C2-epoxy-1%2C2-dihydrophenyl)acetyl-CoA isomerase PaaG | -2.75 |
| **OB946_05255** | DUF3015 family protein | -2.81 |
| **OB946_09260** | zonular occludens toxin domain-containing protein | -2.88 |
| **OB946_10590** | fimbrial protein | -2.89 |
| **OB946_04785** | type I-F CRISPR-associated endoribonuclease Cas6/Csy4 | -2.93 |
| **OB946_00365** | zinc metallochaperone GTPase ZigA | -2.94 |
| **OB946_04780** | type I-F CRISPR-associated protein Csy3 | -2.95 |
| **OB946_09660** | FAD/NAD(P)-binding oxidoreductase | -3.04 |
| **OB946_03140** | Hpt domain-containing protein | -3.10 |
| **OB946_09670** | MBL fold metallo-hydrolase | -3.17 |
| **OB946_05465** | hypothetical protein | -3.23 |
| **OB946_08120** | PEGA domain-containing protein | -3.26 |
| **OB946_09425** | isochorismatase family protein | -3.29 |
| **OB946_11460** | ATP-binding cassette domain-containing protein | -3.32 |
| **OB946_01570** | hypothetical protein | -3.32 |
| **OB946_14630** | PilT/PilU family type 4a pilus ATPase | -3.37 |
| **OB946_08640** | catechol 1%2C2-dioxygenase | -3.46 |
| **OB946_11655** | DapH/DapD/GlmU-related protein | -3.52 |
| **OB946_07645** | acetyl-CoA C-acetyltransferase | -3.52 |
| **OB946_00200** | MFS transporter | -3.54 |
| **OB946_15000** | peptidoglycan-binding protein LysM | -3.65 |
| **OB946_08645** | muconolactone Delta-isomerase | -3.74 |
| **OB946_11665** | phenylacetate--CoA ligase PaaK | -3.76 |
| **OB946_09280** | hypothetical protein | -4.01 |
| **OB946_08785** | hydrolase | -4.32 |
| **OB946_09205** | hypothetical protein | -4.46 |
| **OB946_10625** | hypothetical protein | -4.57 |
| **OB946_08780** | putative quinol monooxygenase | -4.72 |
| **OB946_11675** | 3-hydroxyacyl-CoA dehydrogenase | -4.73 |
| **OB946_09665** | | -5.13 |
| **OB946_07585** | universal stress protein | -5.16 |

| | | |
|---|---|---|
| **OB946_11670** | 3-oxoadipyl-CoA thiolase | -6.25 |
| **OB946_10620** | DMT family transporter | -7.32 |
| **OB946_01505** | pilin | -9.07 |
| **OB946_09200** | major capsid protein | -14.46 |
| **OB946_09275** | major capsid protein | -14.52 |

a| Fold change cutoff: 2-fold *P* value < 0.01. Differential expression was calculated with DESeq2.

**Supplemental data set 3: Differentially expressed genes in ARC6851 in colistin + diclofenac treatment vs diclofenac.**

| Accession | Annotated gene | Fold change[a] |
|---|---|---|
| **OB946_12790** | SDR family oxidoreductase | 11.54 |
| **OB946_12785** | NAD(P)/FAD-dependent oxidoreductase | 10.04 |
| **OB946_12795** | alpha/beta hydrolase | 5.22 |
| **OB946_00885** | FMN reductase | 4.67 |
| **OB946_09560** | hypothetical protein | 4.64 |
| **OB946_09895** | sulfite exporter TauE/SafE family protein | 4.08 |
| **OB946_01760** | hypothetical protein | 3.97 |
| **OB946_18875** | sulfonate ABC transporter substrate-binding protein | 3.86 |
| **OB946_15220** | TetR/AcrR family transcriptional regulator | 3.72 |
| **OB946_18890** | aliphatic sulfonate ABC transporter permease SsuC | 3.35 |
| **OB946_00880** | dimethyl sulfone monooxygenase SfnG | 3.23 |
| **OB946_18865** | RcnB family protein | 3.07 |
| **OB946_04880** | substrate-binding domain-containing protein | 2.92 |
| **OB946_12420** | transposase | 2.88 |
| **OB946_15495** | substrate-binding domain-containing protein | 2.82 |
| **OB946_10965** | monooxygenase | 2.77 |
| **OB946_15450** | DUF1852 domain-containing protein | 2.77 |
| **OB946_07300** | hypothetical protein | 2.66 |
| **OB946_15490** | MFS transporter | 2.65 |
| **OB946_17300** | hypothetical protein | 2.63 |
| **OB946_16045** | RidA family protein | 2.58 |
| **OB946_04425** | sulfate ABC transporter substrate-binding protein | 2.56 |
| **OB946_04895** | phosphate ABC transporter ATP-binding protein PstB | 2.54 |
| **OB946_13200** | LysE/ArgO family amino acid transporter | 2.52 |

| | | |
|---|---|---|
| **OB946_02760** | RcnB family protein | 2.52 |
| **OB946_12390** | hypothetical protein | 2.46 |
| **OB946_18880** | sulfonate ABC transporter substrate-binding protein | 2.46 |
| **OB946_09460** | anthranilate 1%2C2-dioxygenase small subunit | 2.44 |
| **OB946_09715** | lipoyl synthase | 2.37 |
| **OB946_03085** | DUF488 domain-containing protein | 2.37 |
| **OB946_02780** | hypothetical protein | 2.37 |
| **OB946_13205** | cation diffusion facilitator family transporter | 2.36 |
| **OB946_09940** | GNAT family N-acetyltransferase | 2.36 |
| **OB946_07435** | hypothetical protein | 2.35 |
| **OB946_18255** | hypothetical protein | 2.35 |
| **OB946_12440** | hypothetical protein | 2.32 |
| **OB946_18800** | hypothetical protein | 2.32 |
| **OB946_09500** | MFS transporter | 2.29 |
| **OB946_17865** | site-specific integrase | 2.28 |
| **OB946_02885** | peroxiredoxin | 2.27 |
| **OB946_06760** | hypothetical protein | 2.26 |
| **OB946_03965** | transglycosylase SLT domain-containing protein | 2.21 |
| **OB946_10945** | taurine ABC transporter substrate-binding protein | 2.19 |
| **OB946_13800** | BCCT family transporter | 2.18 |
| **OB946_18985** | hypothetical protein | 2.17 |
| **OB946_04890** | phosphate ABC transporter permease PstA | 2.16 |
| **OB946_03935** | multidrug efflux RND transporter permease subunit | 2.15 |
| **OB946_08435** | 3-carboxy-cis%2Ccis-muconate cycloisomerase | 2.12 |
| **OB946_10395** | hypothetical protein | 2.11 |
| **OB946_15410** | hypothetical protein | 2.11 |
| **OB946_06850** | | 2.10 |
| **OB946_13430** | sulfate adenylyltransferase subunit CysD | 2.10 |
| **OB946_09455** | anthranilate 1%2C2-dioxygenase large subunit | 2.10 |
| **OB946_07115** | hypothetical protein | 2.09 |

| | | |
|---|---|---|
| **OB946_18885** | FMNH2-dependent alkanesulfonate monooxygenase | 2.09 |
| **OB946_04885** | phosphate ABC transporter permease subunit PstC | 2.09 |
| **OB946_00595** | lysozyme inhibitor LprI family protein | 2.08 |
| **OB946_12445** | LysE family transporter | 2.07 |
| **OB946_17670** | signal peptidase II | 2.07 |
| **OB946_18860** | RcnB family protein | 2.05 |
| **OB946_02740** | DUF4184 family protein | 2.05 |
| **OB946_18895** | ATP-binding cassette domain-containing protein | 2.05 |
| **OB946_00240** | hypothetical protein | 2.04 |
| **OB946_18980** | hypothetical protein | 2.04 |
| **OB946_09465** | anthranilate 1%2C2-dioxygenase electron transfer component AntC | 2.04 |
| **OB946_11020** | biotin-independent malonate decarboxylase subunit gamma | 2.04 |
| **OB946_13210** | MerR family transcriptional regulator | 2.02 |
| **OB946_12205** | site-specific integrase | 2.01 |
| **OB946_02160** | membrane protein | 2.01 |
| **OB946_12830** | MarR family transcriptional regulator | -2.01 |
| **OB946_19440** | | -2.02 |
| **OB946_10780** | tautomerase family protein | -2.03 |
| **OB946_19240** | | -2.03 |
| **OB946_01575** | VWA domain-containing protein | -2.03 |
| **OB946_15360** | hypothetical protein | -2.04 |
| **OB946_11975** | divalent metal cation transporter | -2.04 |
| **OB946_04870** | thiamine pyrophosphate-binding protein | -2.04 |
| **OB946_13975** | hypothetical protein | -2.05 |
| **OB946_18025** | TorF family putative porin | -2.06 |
| **OB946_08960** | phosphoribosyltransferase family protein | -2.07 |
| **OB946_06835** | acyl-CoA desaturase | -2.09 |
| **OB946_05255** | DUF3015 family protein | -2.09 |
| **OB946_11660** | phenylacetic acid degradation operon negative regulatory protein PaaX | -2.10 |
| **OB946_08205** | hypothetical protein | -2.12 |
| **OB946_11390** | 3-hydroxyacyl-CoA dehydrogenase | -2.13 |

| | | |
|---|---|---|
| **OB946_03135** | methyl-accepting chemotaxis protein | -2.13 |
| **OB946_00385** | urocanate hydratase | -2.13 |
| **OB946_04045** | formate dehydrogenase accessory sulfurtransferase FdhD | -2.14 |
| **OB946_15250** | mechanosensitive ion channel | -2.16 |
| **OB946_03495** | hypothetical protein | -2.16 |
| **OB946_01450** | pilus assembly protein PilP | -2.16 |
| **OB946_08035** | YcgJ family protein | -2.17 |
| **OB946_07615** | benzoate/H(+) symporter BenE family transporter | -2.17 |
| **OB946_17940** | baseplate J/gp47 family protein | -2.17 |
| **OB946_07685** | thioesterase family protein | -2.19 |
| **OB946_14725** | threonine export protein RhtC | -2.19 |
| **OB946_05105** | TetR/AcrR family transcriptional regulator | -2.20 |
| **OB946_03145** | hypothetical protein | -2.20 |
| **OB946_07570** | hypothetical protein | -2.21 |
| **OB946_04995** | EAL domain-containing protein | -2.22 |
| **OB946_09285** | hypothetical protein | -2.24 |
| **OB946_09255** | hypothetical protein | -2.25 |
| **OB946_01455** | type IV pilus secretin PilQ family protein | -2.30 |
| **OB946_04335** | multidrug effflux MFS transporter | -2.30 |
| **OB946_09310** | hypothetical protein | -2.31 |
| **OB946_06025** | hypothetical protein | -2.31 |
| **OB946_10590** | fimbrial protein | -2.33 |
| **OB946_09425** | isochorismatase family protein | -2.35 |
| **OB946_00365** | zinc metallochaperone GTPase ZigA | -2.36 |
| **OB946_09225** | hypothetical protein | -2.36 |
| **OB946_08785** | hydrolase | -2.37 |
| **OB946_08690** | hypothetical protein | -2.38 |
| **OB946_08495** | DUF6691 family protein | -2.40 |
| **OB946_16105** | hypothetical protein | -2.41 |
| **OB946_19130** | 50S ribosomal protein L34 | -2.41 |
| **OB946_01565** | PilW family protein | -2.43 |

| | | |
|---|---|---|
| **OB946_11685** | enoyl-CoA hydratase-related protein | -2.46 |
| **OB946_12075** | MFS transporter | -2.48 |
| **OB946_14625** | type IV pilus twitching motility protein PilT | -2.48 |
| **OB946_03685** | | -2.50 |
| **OB946_01570** | hypothetical protein | -2.52 |
| **OB946_11680** | 2-(1%2C2-epoxy-1%2C2-dihydrophenyl)acetyl-CoA isomerase PaaG | -2.54 |
| **OB946_10625** | hypothetical protein | -2.56 |
| **OB946_11650** | PaaI family thioesterase | -2.56 |
| **OB946_05345** | | -2.56 |
| **OB946_08365** | nuclear transport factor 2 family protein | -2.59 |
| **OB946_16150** | DUF4124 domain-containing protein | -2.65 |
| **OB946_04785** | type I-F CRISPR-associated endoribonuclease Cas6/Csy4 | -2.67 |
| **OB946_10865** | chromate transporter | -2.67 |
| **OB946_07645** | acetyl-CoA C-acetyltransferase | -2.67 |
| **OB946_01560** | type IV pilus modification protein PilV | -2.70 |
| **OB946_17855** | hypothetical protein | -2.71 |
| **OB946_07585** | universal stress protein | -2.72 |
| **OB946_04780** | type I-F CRISPR-associated protein Csy3 | -2.81 |
| **OB946_15000** | peptidoglycan-binding protein LysM | -2.83 |
| **OB946_03140** | Hpt domain-containing protein | -2.84 |
| **OB946_11665** | phenylacetate--CoA ligase PaaK | -2.85 |
| **OB946_19385** | | -2.94 |
| **OB946_10620** | DMT family transporter | -3.12 |
| **OB946_05540** | Ish1 domain-containing protein | -3.17 |
| **OB946_09280** | hypothetical protein | -3.21 |
| **OB946_14630** | PilT/PilU family type 4a pilus ATPase | -3.28 |
| **OB946_06180** | hypothetical protein | -3.28 |
| **OB946_01555** | GspH/FimT family pseudopilin | -3.30 |
| **OB946_11655** | DapH/DapD/GlmU-related protein | -3.40 |
| **OB946_09665** | | -3.41 |
| **OB946_11675** | 3-hydroxyacyl-CoA dehydrogenase | -3.42 |

| | | |
|---|---|---|
| **OB946_00200** | MFS transporter | -3.43 |
| **OB946_09205** | hypothetical protein | -4.19 |
| **OB946_11670** | 3-oxoadipyl-CoA thiolase | -4.72 |
| **OB946_01505** | pilin | -5.91 |
| **OB946_09275** | major capsid protein | -9.25 |
| **OB946_09200** | major capsid protein | -9.35 |

a| Fold change cutoff: 2-fold *P* value < 0.01. Differential expression was calculated with DESeq2.

**Supplemental data set 4: Differentially expressed proteins in ARC6851 in colistin + diclofenac treatment vs DMSO.**

| Accession | Annotated protein | Fold change[a] |
|---|---|---|
| UYC77247.1 | hypothetical protein | 9.09 |
| UYC76564.1 | Transcriptional regulator, AcrR family | 8.53 |
| UYC78812.1 | Beta-ketoadipate enol-lactone hydrolase (EC 3.1.1.24) | 5.38 |
| **UYC76578.1** | **NADH-ubiquinone oxidoreductase chain M (EC 1.6.5.3)** | **4.92** |
| **UYC75585.1** | **Putative sulfate permease** | **4.62** |
| UYC77596.1 | hypothetical protein | 4.34 |
| UYC76922.1 | CDP-diacylglycerol--glycerol-3-phosphate 3-phosphatidyltransferase (EC 2.7.8.5) | 3.67 |
| UYC75975.1 | Uncharacterized metal ion transporter YcsG, Mn(2+)/Fe(2+) NRAMP family | 3.59 |
| UYC78867.1 | TetR/AcrR family transcriptional regulator | 3.56 |
| UYC78269.1;UYC78378.1 | minor capsid protein | 3.53 |
| UYC78817.1 | 3-dehydroshikimate dehydratase (EC 4.2.1.118) | 3.17 |
| UYC75736.1 | Glutamate/aspartate ABC transporter, permease protein GltK (TC 3.A.1.3.4) | 3.14 |
| **UYC76566.1** | **Membrane fusion component of MSF-type tripartite multidrug efflux system** | **3.00** |
| UYC78767.1 | hypothetical protein | 2.64 |
| UYC77309.1 | DNA-3-methyladenine glycosylase I | 2.63 |
| UYC77134.1 | DedA protein | 2.58 |
| UYC76190.1 | Arginine exporter protein ArgO | 2.48 |
| UYC77766.1 | Putative transmembrane protein | 2.46 |
| UYC76238.1 | 2-oxoglutarate dehydrogenase complex, dehydrogenase component | 2.43 |
| UYC78596.1 | FIG00352022: hypothetical protein | 2.37 |
| UYC76037.1 | Transcriptional regulator, HxlR family | 2.33 |
| UYC75575.1 | Bis(5'-nucleosyl)-tetraphosphatase (asymmetrical) (EC 3.6.1.17) | 2.32 |
| UYC75915.1 | ATP-dependent helicase | 2.32 |
| UYC76930.1 | hypothetical protein | 2.22 |
| **UYC76504.1** | **Multimeric flavodoxin WrbA** | **2.16** |
| UYC76492.1 | FIG00350262: hypothetical protein | 2.14 |
| **UYC78418.1** | **Probable transcription regulator protein of MDR efflux pump cluster** | **2.08** |
| UYC76035.1 | hypothetical protein | 2.00 |
| **UYC75917.1** | **Acyl-coenzyme A thioesterase PaaD (Pse.pu.) (E. coli PaaI)** | **2.00** |
| UYC78003.1 | D-serine/D-alanine/glycine transporter | -2.00 |

| | | |
|---|---|---|
| UYC76193.1 | FIG00350187: hypothetical protein | -2.01 |
| UYC78169.1 | 5'-nucleotidase SurE (EC 3.1.3.5) | -2.02 |
| UYC79102.1 | Histone acetyltransferase HPA2 and related acetyltransferases | -2.04 |
| UYC77125.1 | Cell division protein | -2.04 |
| UYC76152.1 | Oxidoreductase | -2.04 |
| UYC78598.1 | Acetyltransferase, GNAT family | -2.06 |
| UYC78159.1 | Large repetitive protein | -2.12 |
| UYC77384.1 | Nucleoside-binding outer membrane protein | -2.16 |
| UYC76795.1 | LSU ribosomal protein L33p @ LSU ribosomal protein L33p, zinc-independent | -2.16 |
| UYC75868.1 | 3-hydroxyacyl-CoA dehydrogenase | -2.18 |
| UYC77509.1 | hypothetical protein OB946_01300 | -2.20 |
| UYC77005.1 | hypothetical protein | -2.22 |
| UYC76852.1 | Type III effector HopPmaJ | -2.24 |
| UYC76907.1 | Leader peptidase (Prepilin peptidase) (EC 3.4.23.43) / N-methyltransferase (EC 2.1.1.-) | -2.25 |
| UYC76219.1 | Urease accessory protein UreE | -2.25 |
| UYC77374.1 | hypothetical protein OB946_00585 | -2.26 |
| **UYC76455.1** | **Type IV pilus assembly ATPase component PilU** | **-2.27** |
| **UYC77535.1** | **Type IV pilus biogenesis protein PilQ** | **-2.31** |
| UYC76020.1 | hypothetical protein OB946_12265 | -2.33 |
| UYC79027.1 | hypothetical protein OB946_02540 | -2.34 |
| UYC77207.1 | 1,6-anhydro-N-acetylmuramyl-L-alanine amidase | -2.38 |
| UYC78379.1 | hypothetical protein OB946_06090 | -2.44 |
| UYC75698.1 | hypothetical protein | -2.53 |
| UYC76183.1 | FIG00350110: hypothetical protein | -2.55 |
| UYC76926.1 | hypothetical protein OB946_06090 | -2.57 |
| **UYC77836.1** | **Twitching motility protein PilH** | **-2.57** |
| UYC79047.1 | hypothetical protein OB946_17215 | -2.58 |
| **UYC79020.1** | **Type IV pilus biogenesis protein PilP** | **-2.60** |
| UYC76816.1 | hypothetical protein | -2.61 |
| UYC78114.1 | FKBP-type peptidyl-prolyl cis-trans isomerase SlyD (EC 5.2.1.8) | -2.63 |
| **UYC77534.1** | **Type IV pilus biogenesis protein PilO** | **-2.69** |
| UYC76426.1 | YfdQ family protein | -2.71 |
| UYC76450.1 | OsmC/Ohr family protein | -2.75 |
| **UYC77533.1** | **Type IV pilus biogenesis protein PilN** | **-2.84** |
| UYC76525.1 | LSU ribosomal protein L32p @ LSU ribosomal protein L32p, zinc-independent | -2.91 |
| **UYC77532.1** | **Type IV pilus biogenesis protein PilM** | **-2.98** |
| UYC78428.1 | Type II secretory pathway, ATPase PulE/Tfp pilus assembly pathway, ATPase PilB | -3.00 |

| | | |
|---|---|---|
| UYC75926.1 | 1,2-phenylacetyl-CoA epoxidase, subunit D (EC 1.14.13.149) | -3.02 |
| UYC77644.1 | SSU ribosomal protein S19p (S15e) | -3.07 |
| **UYC77835.1** | **Twitching motility protein PilG** | **-3.11** |
| UYC76608.1 | hypothetical protein | -3.14 |
| UYC76488.1 | Protoporphyrinogen IX oxidase, novel form, HemJ (EC 1.3.-.-) | -3.20 |
| UYC76441.1 | endonuclease/exonuclease/phosphatase family protein | -3.24 |
| UYC77519.1 | Uncharacterized amino acid permease, GabP family | -3.27 |
| UYC78424.1 | Uncharacterized protease YegQ | -3.31 |
| **UYC76906.1** | **Type IV fimbrial assembly protein PilC** | **-3.68** |
| UYC77157.1 | D-serine/D-alanine/glycine transporter | -3.79 |
| UYC75670.1 | MotA/TolQ/ExbB proton channel family protein | -3.83 |
| UYC75523.1 | Aspartate ammonia-lyase (EC 4.3.1.1) | -4.20 |
| UYC78695.1 | GTP 3',8-cyclase (EC 4.1.99.22) | -4.66 |
| UYC79070.1 | PEGA domain-containing protein | -4.74 |
| UYC77025.1 | BRCT domain-containing protein | -5.19 |
| UYC78168.1 | hypothetical protein | -5.22 |
| UYC78954.1;UYC78945.1 | hypothetical protein OB946_09210 | -5.46 |
| UYC77840.1 | hypothetical protein | -6.10 |
| **UYC77837.1** | **Type IV pili signal transduction protein PilI** | **-6.70** |
| **UYC77838.1** | **Type IV pilus biogenesis protein PilJ** | -6.86 |
| UYC78963.1;UYC78949.1 | zonular occludens toxin domain-containing protein | -7.16 |
| **UYC77558.1** | **Type IV fimbrial biogenesis protein PilY1** | **-7.90** |
| UYC76460.1 | Hemerythrin domain protein | -10.26 |
| **UYC77545.1** | **Type IV pilin PilA** | **-10.95** |
| **UYC77839.1** | **Twitching motility protein PilG** | **-18.79** |
| UYC76989.1 | Helix-turn-helix, Fis-type | -22.74 |

**Supplemental data set 5: Differentially expressed proteins in ARC6851 in colistin + diclofenac treatment vs colistin.**

| Accession | Annotated protein | Fold change[a] |
|---|---|---|
| UYC76564.1 | Transcriptional regulator, AcrR family | 8.84 |
| UYC79064.1 | Glycine-rich cell wall structural protein precursor | 5.42 |
| UYC76804.1 | hypothetical protein | 4.88 |
| UYC77596.1 | hypothetical protein | 4.18 |
| UYC78867.1 | TetR/AcrR family transcriptional regulator | 3.27 |
| **UYC76566.1** | **Membrane fusion component of MSF-type tripartite multidrug efflux system** | **2.88** |
| UYC76190.1 | Arginine exporter protein ArgO | 2.85 |
| UYC75585.1 | Putative sulfate permease | 2.75 |
| UYC79067.1 | Mg(2+) transport ATPase, P-type (EC 3.6.3.2) | 2.67 |
| UYC77766.1 | Putative transmembrane protein | 2.63 |
| UYC76922.1 | CDP-diacylglycerol--glycerol-3-phosphate 3-phosphatidyltransferase (EC 2.7.8.5) | 2.62 |
| UYC77134.1 | DedA protein | 2.61 |
| UYC77594.1 | hypothetical protein | 2.37 |
| UYC78127.1 | hypothetical protein | 2.19 |
| UYC78814.1 | 4-carboxymuconolactone decarboxylase (EC 4.1.1.44) | 2.09 |
| UYC78830.1;UYC76067.1 | Outer membrane low permeability porin, OprD family | 2.07 |
| UYC77224.1 | Alkanesulfonate ABC transporter substrate-binding protein SsuA | 2.06 |
| UYC75575.1 | Bis(5'-nucleosyl)-tetraphosphatase (asymmetrical) (EC 3.6.1.17) | 2.04 |
| UYC78809.1 | 3-oxoadipate CoA-transferase subunit B (EC 2.8.3.6) | 2.01 |
| UYC78708.1 | FIG00350630: hypothetical protein | 2.01 |
| UYC77612.1 | hypothetical protein | -1.99 |
| UYC76332.1 | cupin domain-containing protein | -2.00 |
| UYC77740.1 | probable membrane protein STY1534 | -2.01 |
| UYC79271.1 | sel1 repeat family protein | -2.01 |
| UYC78765.1 | hypothetical protein | -2.02 |
| UYC76464.1 | Phosphoglucomutase (EC 5.4.2.2) @ Phosphomannomutase (EC 5.4.2.8) | -2.03 |
| UYC78171.1 | Transcriptional regulator, LysR family | -2.04 |
| UYC77687.1 | TonB, C-terminal | -2.06 |
| UYC78135.1 | type I-F CRISPR-associated protein Csy3 | -2.06 |
| UYC78409.1 | DUF4760 domain-containing protein | -2.09 |
| UYC76677.1 | hypothetical protein OB946_15875 | -2.11 |
| UYC78968.1;UYC78957.1 | hypothetical protein OB946_09285 | -2.11 |
| UYC75878.1 | Phosphonoacetaldehyde hydrolase (EC 3.11.1.1) | -2.17 |
| UYC76653.1 | Ribosome hibernation promoting factor Hpf | -2.21 |
| UYC77193.1 | hypothetical protein OB946_18720 | -2.22 |
| UYC75967.1 | hypothetical protein OB946_11930 | -2.23 |

| | | |
|---|---|---|
| UYC75711.1 | Crossover junction endodeoxyribonuclease RuvC (EC 3.1.22.4) | -2.23 |
| UYC77434.1 | hypothetical protein | -2.23 |
| UYC76495.1 | Carnitine monooxygenase, oxygenase component CntA | -2.24 |
| UYC76155.1 | AAA domain-containing protein | -2.25 |
| UYC76058.1 | Cu(I)-responsive transcriptional regulator | -2.27 |
| UYC77509.1 | hypothetical protein OB946_01300 | -2.27 |
| UYC77414.1 | MBL-fold metallo-hydrolase superfamily | -2.29 |
| UYC75947.1 | hypothetical protein | -2.33 |
| **UYC76454.1** | **Twitching motility protein PilT** | **-2.34** |
| UYC75562.1 | hypothetical protein | -2.36 |
| UYC75922.1 | 3-hydroxyadipyl-CoA dehydrogenase | -2.37 |
| UYC78877.1 | LysR substrate-binding domain-containing protein | -2.38 |
| UYC78632.1 | Periplasmic chorismate mutase I precursor (EC 5.4.99.5) | -2.39 |
| UYC76852.1 | Type III effector HopPmaJ | -2.45 |
| UYC78144.1 | Linoleoyl-CoA desaturase (EC 1.14.19.3) | -2.47 |
| UYC77065.1 | FIGfam050825 | -2.52 |
| UYC78923.1 | DEAD/DEAH box helicase family protein | -2.52 |
| UYC77876.1 | Fatty acid desaturase | -2.53 |
| UYC78598.1 | Acetyltransferase, GNAT family | -2.54 |
| UYC77682.1 | Inner membrane protein YihY, formerly thought to be RNase BN | -2.57 |
| **UYC75925.1** | **1,2-phenylacetyl-CoA epoxidase, subunit E (EC 1.14.13.149)** | **-2.60** |
| UYC76139.1 | Probable coniferyl aldehyde dehydrogenase (EC 1.2.1.68);Aldehyde dehydrogenase (EC 1.2.1.3) | -2.61 |
| UYC78143.1 | Flavodoxin reductases (ferredoxin-NADPH reductases) family 1 | -2.66 |
| UYC78133.1 | type I-F CRISPR-associated protein Csy1 | -2.68 |
| UYC78245.1 | hypothetical protein OB946_05385 | -2.68 |
| UYC75633.1 | ABC transporter ATP-binding protein | -2.71 |
| **UYC75920.1** | **Phenylacetate-coenzyme A ligase (EC 6.2.1.30)** | **-2.74** |
| **UYC77836.1** | **Twitching motility protein PilH** | **-2.74** |
| UYC78134.1 | type I-F CRISPR-associated protein Csy2 | -2.77 |
| UYC76907.1 | Leader peptidase (Prepilin peptidase) (EC 3.4.23.43) / N-methyltransferase (EC 2.1.1.-) | -2.78 |
| **UYC77535.1** | **Type IV pilus biogenesis protein PilQ** | **-2.80** |
| **UYC79020.1** | **Type IV pilus biogenesis protein PilP** | **-2.80** |
| **UYC76455.1** | **Type IV pilus assembly ATPase component PilU** | **-2.81** |
| UYC79102.1 | Histone acetyltransferase HPA2 and related acetyltransferases | -2.84 |
| UYC77374.1 | hypothetical protein OB946_00585 | -2.85 |
| **UYC77557.1** | **Type IV fimbrial biogenesis protein PilX** | **-2.86** |
| UYC77161.1 | D-amino acid dehydrogenase (EC 1.4.99.6) | -2.88 |

| | | |
|---|---|---|
| UYC79027.1 | hypothetical protein OB946_02540 | -2.89 |
| UYC77747.1 | hypothetical protein | -2.89 |
| UYC77456.1 | Gamma-aminobutyrate:alpha-ketoglutarate aminotransferase (EC 2.6.1.19) | -2.89 |
| UYC78159.1 | Large repetitive protein | -2.91 |
| **UYC75927.1** | **1,2-phenylacetyl-CoA epoxidase, subunit C (EC 1.14.13.149)** | **-2.93** |
| UYC76816.1 | hypothetical protein | -2.99 |
| **UYC77534.1** | **Type IV pilus biogenesis protein PilO** | **-3.02** |
| **UYC76926.1** | **hypothetical protein OB946_17215** | **-3.07** |
| UYC76152.1 | Oxidoreductase | -3.08 |
| UYC76183.1 | FIG00350110: hypothetical protein | -3.13 |
| UYC77454.1 | gamma-aminobutyrate (GABA) permease | -3.16 |
| UYC76727.1 | FIG00349950: hypothetical protein | -3.18 |
| UYC76608.1 | hypothetical protein | -3.26 |
| UYC75928.1 | 1,2-phenylacetyl-CoA epoxidase, subunit B (EC 1.14.13.149) | -3.27 |
| **UYC77556.1** | **Type IV fimbrial biogenesis protein PilW** | **-3.28** |
| UYC76219.1 | Urease accessory protein UreE | -3.30 |
| **UYC75929.1** | **1,2-phenylacetyl-CoA epoxidase, subunit A (EC 1.14.13.149)** | **-3.32** |
| UYC79070.1 | PEGA domain-containing protein | -3.45 |
| **UYC77533.1** | **Type IV pilus biogenesis protein PilN** | **-3.49** |
| UYC77835.1 | twitching motility protein PilG | -3.75 |
| **UYC77532.1** | **Type IV pilus biogenesis protein PilM** | **-3.76** |
| UYC75753.1 | Tautomerase | -3.90 |
| UYC77005.1 | hypothetical protein | -3.97 |
| UYC75523.1 | Aspartate ammonia-lyase (EC 4.3.1.1) | -4.33 |
| UYC78428.1 | Type II secretory pathway, ATPase PulE/Tfp pilus assembly pathway, ATPase PilB | -4.51 |
| UYC77340.1 | Formiminoglutamase (EC 3.5.3.8) | -4.90 |
| **UYC76906.1** | **Type IV fimbrial assembly protein PilC** | **-5.17** |
| UYC79047.1 | type I-F CRISPR-associated endoribonuclease Cas6/Csy4 | -5.34 |
| UYC77840.1 | hypothetical protein | -5.70 |
| UYC78379.1 | hypothetical protein OB946_06090 | -5.73 |
| UYC78474.1 | Adenosylmethionine-8-amino-7-oxononanoate aminotransferase (EC 2.6.1.62) | -5.74 |
| UYC78424.1 | Uncharacterized protease YegQ | -6.00 |
| UYC76692.1 | Late competence protein ComEA, DNA receptor | -6.04 |
| UYC77025.1 | BRCT domain-containing protein | -7.07 |
| UYC76860.1 | hypothetical protein | -7.28 |
| UYC78168.1 | hypothetical protein | -7.38 |
| UYC78954.1;UYC78945.1 | hypothetical protein OB946_09210 | -7.79 |
| **UYC77837.1** | **Type IV pili signal transduction protein PilI** | **-8.88** |

| | | |
|---|---|---|
| **UYC77838.1** | **Type IV pilus biogenesis protein PilJ** | **-9.97** |
| UYC75926.1 | 1,2-phenylacetyl-CoA epoxidase, subunit D (EC 1.14.13.149) | -11.68 |
| **UYC77545.1** | **Type IV pilin PilA** | **-13.07** |
| **UYC77558.1** | **Type IV fimbrial biogenesis protein PilY1** | **-13.25** |
| UYC76460.1 | Hemerythrin domain protein | -15.60 |
| UYC76989.1 | Helix-turn-helix, Fis-type | -30.09 |
| UYC78963.1;UYC78949.1 | zonular occludens toxin domain-containing protein | -31.31 |
| **UYC77839.1** | **Twitching motility protein PilG** | **-32.87** |

**a|** Fold change cutoff: 2-fold with a p-value < 0.05. Student's unpaired *t* test.

**Supplemental data set 6: Differentially expressed proteins in ARC6851 in colistin and diclofenac treatment vs diclofenac.**

| Accession | Annotated protein | Fold change[a] |
|---|---|---|
| UYC75722.1 | Transcriptional regulator, LysR family | 24.51 |
| UYC76950.1 | Protein translocase subunit SecE | 11.54 |
| UYC76459.1 | FIG00351543: hypothetical protein | 7.16 |
| UYC76922.1 | CDP-diacylglycerol--glycerol-3-phosphate 3-phosphatidyltransferase (EC 2.7.8.5) | 6.74 |
| UYC77596.1 | hypothetical protein | 6.59 |
| UYC77191.1 | glycosyltransferase family 4 protein | 4.97 |
| UYC76564.1 | Transcriptional regulator, AcrR family | 4.21 |
| UYC78767.1 | hypothetical protein | 3.90 |
| UYC76504.1 | Multimeric flavodoxin WrbA | 3.17 |
| UYC77594.1 | hypothetical protein | 3.12 |
| UYC77597.1 | hypothetical protein | 3.10 |
| UYC75585.1 | Putative sulfate permease | 3.00 |
| UYC78477.1 | Bsr8028 protein | 2.68 |
| **UYC76566.1** | **Membrane fusion component of MSF-type tripartite multidrug efflux system** | **2.65** |
| UYC76053.1 | FIG00351239: hypothetical protein | 2.49 |
| UYC78867.1 | TetR/AcrR family transcriptional regulator | 2.45 |
| UYC75575.1 | Bis(5'-nucleosyl)-tetraphosphatase (asymmetrical) (EC 3.6.1.17) | 2.44 |
| UYC75736.1 | Glutamate/aspartate ABC transporter, permease protein GltK (TC 3.A.1.3.4) | 2.26 |
| UYC76238.1 | 2-oxoglutarate dehydrogenase complex, dehydrogenase component | 2.24 |
| UYC77860.1 | putative signal peptide;hypothetical protein | 2.22 |
| UYC78817.1 | 3-dehydroshikimate dehydratase (EC 4.2.1.118) | 2.11 |
| UYC76321.1 | hypothetical protein OB946_13920 | 2.06 |
| UYC78878.1 | hypothetical protein | 2.06 |
| **UYC75917.1** | **Acyl-coenzyme A thioesterase PaaD (Pse.pu.) (E. coli PaaI)** | **2.02** |
| UYC77509.1 | hypothetical protein OB946_01300 | -2.01 |
| UYC76914.1 | UPF0301 protein YqgE | -2.05 |
| **UYC76455.1** | **Type IV pilus assembly ATPase component PilU** | **-2.07** |
| **UYC77535.1** | **Type IV pilus biogenesis protein PilQ** | **-2.08** |
| UYC77372.1 | VgrG protein | -2.08 |
| UYC77374.1 | hypothetical protein OB946_00585 | -2.09 |
| UYC76152.1 | Oxidoreductase | -2.10 |
| UYC77005.1 | hypothetical protein | -2.10 |
| UYC76020.1 | hypothetical protein OB946_12265 | -2.12 |
| UYC76727.1 | FIG00349950: hypothetical protein | -2.12 |

| | | |
|---|---|---|
| UYC75717.1;UYC78972.1 | Fimbrial protein precursor | -2.16 |
| UYC79027.1 | hypothetical protein OB946_02540 | -2.19 |
| UYC77207.1 | 1,6-anhydro-N-acetylmuramyl-L-alanine amidase | -2.22 |
| UYC76525.1 | LSU ribosomal protein L32p @ LSU ribosomal protein L32p, zinc-independent | -2.24 |
| UYC76495.1 | Carnitine monooxygenase, oxygenase component CntA | -2.27 |
| **UYC77556.1** | **Type IV fimbrial biogenesis protein PilW** | **-2.28** |
| UYC79047.1 | type I-F CRISPR-associated endoribonuclease Cas6/Csy4 | -2.34 |
| **UYC79020.1** | **Type IV pilus biogenesis protein PilP** | **-2.39** |
| UYC76816.1 | hypothetical protein | -2.40 |
| UYC77384.1 | Nucleoside-binding outer membrane protein | -2.43 |
| **UYC77534.1** | **Type IV pilus biogenesis protein PilO** | **-2.47** |
| **UYC77533.1** | **Type IV pilus biogenesis protein PilN** | **-2.48** |
| UYC76450.1 | OsmC/Ohr family protein | -2.51 |
| UYC76183.1 | FIG00350110: hypothetical protein | -2.56 |
| UYC77644.1 | SSU ribosomal protein S19p (S15e) | -2.63 |
| UYC76907.1 | Leader peptidase (Prepilin peptidase) (EC 3.4.23.43) / N-methyltransferase (EC 2.1.1.-) | -2.64 |
| UYC76426.1 | YfdQ family protein | -2.64 |
| **UYC77835.1** | **Twitching motility protein PilG** | **-2.67** |
| UYC77519.1 | Uncharacterized amino acid permease, GabP family | -2.70 |
| UYC76926.1 | hypothetical protein OB946_17215 | -2.73 |
| UYC78424.1 | Uncharacterized protease YegQ | -2.75 |
| UYC78428.1 | Type II secretory pathway, ATPase PulE/Tfp pilus assembly pathway, ATPase PilB | -2.77 |
| UYC76608.1 | hypothetical protein | -2.88 |
| UYC76692.1 | Late competence protein ComEA, DNA receptor | -2.94 |
| UYC75670.1 | MotA/TolQ/ExbB proton channel family protein | -3.01 |
| **UYC77532.1** | **Type IV pilus biogenesis protein PilM** | **-3.03** |
| UYC78382.1;UYC78272.1 | hypothetical protein | -3.20 |
| UYC75750.1 | hypothetical protein | -3.38 |
| UYC78168.1 | hypothetical protein | -3.45 |
| **UYC76906.1** | **Type IV fimbrial assembly protein PilC** | **-3.46** |
| UYC76219.1 | Urease accessory protein UreE | -3.47 |
| UYC76229.1 | Citrate/H+ symporter of CitMHS family | -3.55 |
| UYC75523.1 | Aspartate ammonia-lyase (EC 4.3.1.1) | -4.13 |
| UYC79070.1 | PEGA domain-containing protein | -4.43 |
| UYC77157.1 | D-serine/D-alanine/glycine transporter | -4.54 |
| UYC77025.1 | BRCT domain-containing protein | -4.77 |
| UYC79060.1 | Uncharacterized UPF0033 protein | -4.77 |
| UYC76860.1 | hypothetical protein | -4.96 |
| UYC78954.1;UYC78945.1 | hypothetical protein OB946_09210 | -5.76 |

| | | |
|---|---|---|
| **UYC77837.1** | **type IV pili signal transduction protein PilI** | **-5.94** |
| **UYC77838.1** | **Type IV pilus biogenesis protein PilJ** | **-6.04** |
| UYC76488.1 | Protoporphyrinogen IX oxidase, novel form, HemJ (EC 1.3.-.-) | -6.61 |
| **UYC77558.1** | **Type IV fimbrial biogenesis protein PilY1** | **-6.73** |
| UYC76460.1 | Hemerythrin domain protein | -7.16 |
| **UYC77545.1** | **Type IV pilin PilA** | **-9.36** |
| **UYC77839.1** | **Twitching motility protein PilG** | **-17.83** |
| UYC76989.1 | Helix-turn-helix, Fis-type | -18.10 |
| UYC78963.1;UYC78949.1 | zonular occludens toxin domain-containing protein | -23.91 |

**a|** Fold change cutoff: 2-fold with a p-value < 0.05. Student's unpaired *t* test.
